# Prediction-based highly sensitive CRISPR off-target validation using target-specific DNA enrichment

**DOI:** 10.1101/2019.12.31.889626

**Authors:** Seung-Hun Kang, Wi-jae Lee, Ju-Hyun An, Jong-Hee Lee, Young-Hyun Kim, Hanseop Kim, Yeounsun Oh, Young-Ho Park, Yeung Bae Jin, Bong-Hyun Jun, Junho K Hur, Sun-Uk Kim, Seung Hwan Lee

**Affiliations:** Futuristic Animal Resource & Research Center (FARRC), Korea Research Institute of Bioscience and Biotechnology (KRIBB), Cheongju, Korea; National Primate Research Center (NPRC), Korea Research Institute of Bioscience and Biotechnology (KRIBB), Cheongju, Korea; Department of Functional Genomics, KRIBB School of Bioscience, Korea University of Science and Technology (UST), Daejeon, Korea; Department of Pathology, College of Medicine, Kyung Hee University, Seoul 02447, Republic of Korea; Department of Biomedical Science, Graduate School, Kyung Hee University, Seoul 02447, Republic of Korea; Department of Medicine, Graduate School, Kyung Hee University, Seoul 02447, Republic of Korea; School of Life Sciences and Biotechnology, BK21 Plus KNU Creative BioResearch Group, Kyungpook National University, Daegu, Republic of Korea; Department of Bioscience and Biotechnology, Konkuk University, Seoul 143-701, Korea; Division of Biotechnology, College of Life Science and Biotechnology, Korea University, Seoul 02841, Republic of Korea

## Abstract

CRISPR effectors, which comprise a CRISPR-Cas protein and a guide (g)RNA derived from the bacterial immune system, are widely used to induce double-strand breaks in target DNA and activate the *in-vivo* DNA repair system for target-specific genome editing. When the gRNA recognizes genomic loci with sequences that are similar to the target, deleterious and often carcinogenic mutations can occur. Off-target mutations with a frequency below 0.5% remain mostly undetected by current genome-wide off-target detection techniques. In this study, we developed a method to effectively detect extremely small amounts of mutated DNA based on predicted off-target-specific amplification. We used various genome editors, including CRISPR-Cpf1, Cas9, and an adenine base editor, to induce intracellular genome mutations. The CRISPR amplification method detected off-target mutations at a significantly higher rate (1.6∼984 fold increase) than did an existing targeted amplicon sequencing method. In the near future, CRISPR amplification in combination with genome-wide off-target detection methods will allow to detect genome editor-induced off-target mutations with high sensitivity and in a non-biased manner.

## Introduction

The CRISPR-Cas system, consisting of various Cas proteins and guide (g) RNA, is a bacterial or archaea immune system required to defend viral DNA through target sequence specific cleavage. The CRISPR-Cas system allows targeted editing of genes of interest in various organisms, from bacteria to humans, and has versatile applications *in vivo*^1, 2^. However, when the CRISPR system recognizes sequences similar to the target sequence, deleterious off-target mutations can occur, especially in mammals given their large genomes, which can lead to malfunctions^3^. Off-targeting issues due to guide (g)RNA characteristics have been reported for various CRISPR effectors, such as Cas9 and Cas12 (Cpf1)^4-8^. Consistently, there is a need for precise control of CRISPR-Cas function, and in particular, for a method to develop CRISPR effectors^9-11^ that precisely target a desired locus while avoiding off-target effects, before CRISPR effectors can be introduced for human therapeutic purposes^12^.

To date, various methods, mostly based on next-generation sequencing (NGS), have been developed to detect the off-target mutations *in vivo*^13^. These methods detect genome-wide off-target mutations in an unbiased fashion both inside and outside the cell. However, the current methods often result in ambiguous or no detection of off-target mutations, particularly, when they are below the sequencing error rate (<0.5%). In addition, these methods detect non-common variations other than shared off-target mutations, and thus require additional validation. In order to measure a small amount of off-target mutations caused by the above effectors with high sensitivity, a method to enrich and relatively amplify the mutated over the much more abundant wild-type DNA using CRISPR endonucleases based on previous method(<u>C</u>RISPR-mediated, <u>u</u>ltrasensitive detection of target DNA (CUT)-PCR) was developed^14^. First, all off-target candidate sequences similar to the given CRISPR effector target sequence are selected by *in-silico* prediction. Subsequently, wild-type DNA that does not have mutations in each of the off-target candidate sites is eliminated by the effector to enrich mutant DNA, which is then PCR-amplified, thus enabling the detection of extremely small amounts of mutant DNA with a sensitivity superior to that of conventional deep sequencing methods. CRISPR amplification based off-target mutation enrichment was never tried with genomic DNA before and we further enhanced this method with direct amplification on cell extracted genomic DNA.

With the rapid advancements in CRISPR-Cas genome editing technologies, their application in human gene therapy is being considered^1, 15^. Therefore, there is an urgent need for a method to accurately detect whether or not CRISPR effectors operate at unwanted off-target sites. This study aimed to develop a method for detecting very small amounts of off-target mutations (below the detection limit of deep sequencing) derived from genetic modification of specific sequences using CRISPR amplification technology *in vivo* with high accuracy.

## Results

### Effective mutant DNA fragment enrichment with CRISPR amplification

The existing NGS methods have limited sensitivity in detecting genome-wide off-target mutations induced by CRISPR effectors *in vivo*. To overcome this limitation, we developed a method for amplifying and analyzing a small amount of mutations on CRISPR effector treated genomic DNA. To detect the off-target mutations, we designed a CRISPR amplification method to allow relative amplification of a small amount of mutant DNA fragments over wild-type DNA. The principle of this method is shown in **(Fig. 1a)**. First, the off-target sequences similar to the target sequence are predicted *in silico*. Then, genomic DNA is extracted from CRISPR-edited cells, and on-target and predicted off-target genome sequences are PCR-amplified.

Subsequently, the amplicons are processed by the CRISPR effector (using the same or optimally designed gRNA to remove wild-type DNA other than mutant DNA), and the enriched mutant DNA fragments are PCR-amplified. Amplicons obtained by 3 times repeated CRISPR effector cutting and PCR amplification are then barcoded by nested PCR and subjected to NGS. The indel frequency (%) is calculated to evaluate the presence of off-target mutations. To verify that mutated DNA was indeed relatively amplified over non-mutated DNA by this method, we edited HEK293FT cells using the CRISPR-Cpf1 system to induce indels in target sequences and applied the CRISPR amplification method to genomic DNA samples extracted from the cells **(Fig. 1b–d)**. As shown in **(Fig. 1b)**, targeted amplicon sequencing and genotyping**(Supplementary Fig. 1a)** enabled the detection of extremely low concentrations of mutant DNA fragments (∼1/100,000%) through multiple rounds of amplification by wild-type DNA specific cleavage**(Fig. 1c, Top)**. Only one cycle of CRISPR amplification was used to detect mutant genes with the rate around 0.01%, and three repeated assays allowed determining the indel mutation rate as low as 0.00001% **(Fig. 1c, Bottom)**. When analyzing the pattern of mutations that were relatively amplified by CRISPR amplification up to three times for each diluted sample from original CRISPR effector treated genomic DNA, deletion patterns were observed mainly in the CRISPR-Cpf1 target sequence, in line with previous reports^16,17^ **(Fig. 1d)**. When the wild-type DNA was removed by the CRISPR effector, DNA fragments containing deletion-type mutations **(Fig. 1d, Right)** of various sizes were effectively amplified over non-mutated DNA fragments. The deletion pattern for the CRISPR-Cpf1 effector revealed that DNA fragments with large deletions tended to be more effectively amplified than fragments with smaller deletions **(Fig. 1d, Left)**. The results indicated that a 1-bp alteration from the original sequence within the protospacer region leads to a high probability of re-cleavage by the CRISPR effector, and mutated DNA with larger indels is relatively better amplified.

**Fig. 1.**
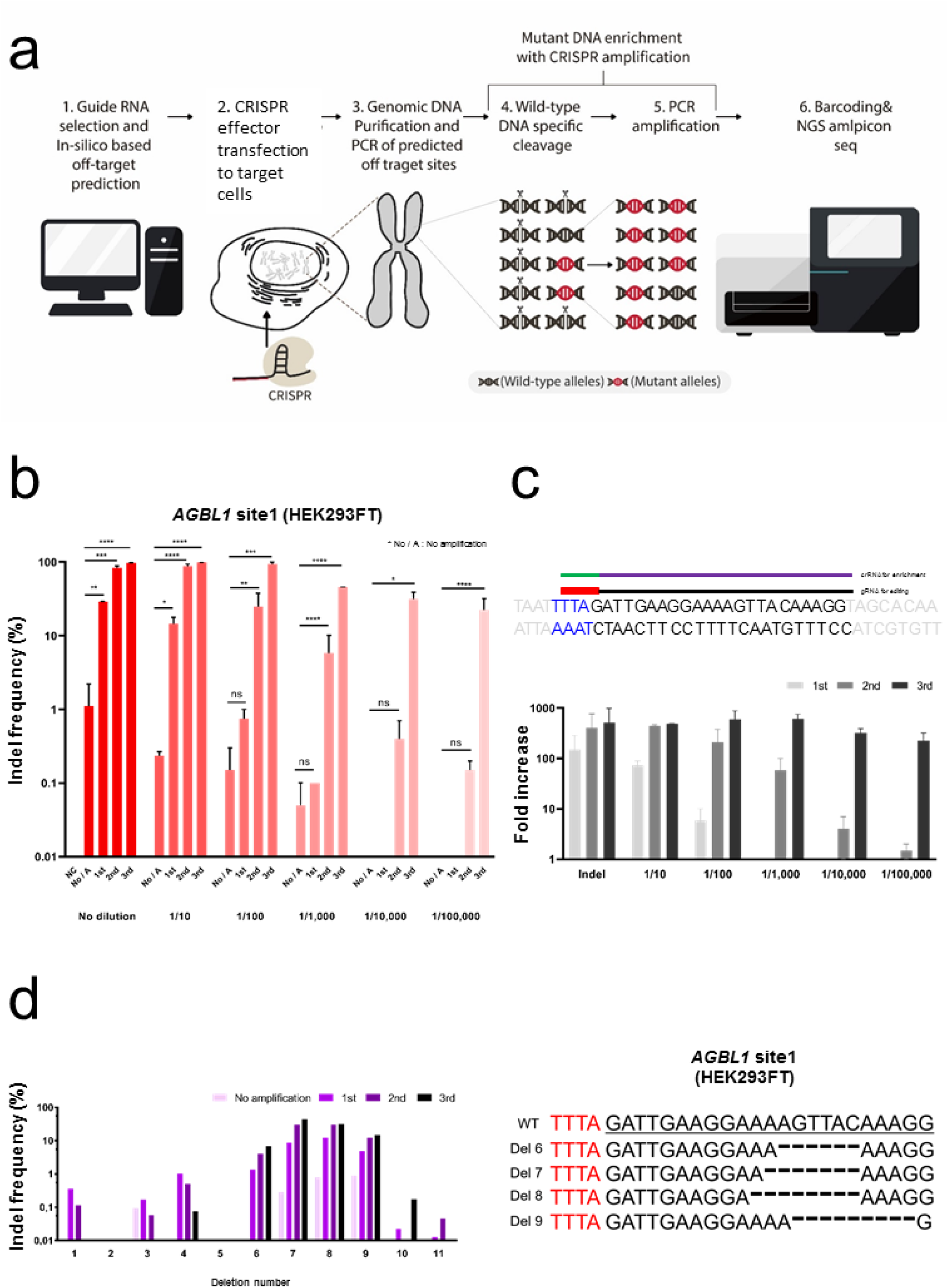
Schematic representation of the detection of off-target mutations using the CRISPR amplification method developed in this study. **(a)** Workflow of the off-target detection method using CRISPR amplification. (1) Based on candidate off-target sequences predicted *in silico*, a gRNA is designed to enrich off-target sequences by subsequent CRISPR cleavage. (2) The CRISPR effector is transfected into cells to induce specific mutations. (3) Genomic DNA is extracted from the cells and predicted off-target loci are PCR-amplified. (4) The PCR amplicons are cleaved by the CRISPR effector. (5) DNA with non-cleaved indel mutations are amplified preferentially over wild-type DNA. (6) The amplified DNA is barcoded and analyzed by NGS. **(b)** Quantitative analysis of mutant DNA enrichment by CRISPR amplification. Genomic DNA samples with mutations induced in the target sequence were serially diluted (up to 1/100,000) and amplified. The indel efficiency (%) was calculated by sequencing DNA amplicons obtained from genomic DNA extracted from CRISPR-Cpf1-edited cells. The X-axis represents the degree of dilution of the genomic DNA samples, and the Y-axis represents the indel detection rate (%). *P*-values are calculated using a Tukey’s test(ns: not significant, *P*\*:<0.0332, *P*\*\*:<0.0021, *P*\*\*\*:<0.0002, *P*\*\*\*\*:<0.0001) **(c)** Top: Design of the crRNA for target specific cleavage and mutant DNA enrichment, respectively. crRNA was used to induce target genomic locus mutation by AsCpf1 effector and exactly same crRNA was used for mutant DNA enrichment by CRISPR amplification. Bottom: Fold increase of indel frequency by CRISPR amplification for each diluted samples from originally mutation induced genomic DNA. **(d)** NGS analysis of AsCpf1 induced indel patterns enriched by CRISPR amplification. Left: Gradual amplification(no amplification, primary, secondary and tertiary amplification) of mutant DNAs with different deletion sizes. Each amplification stage is indicated by different colors. Right: Representative enriched mutation patterns from third round CRISPR amplification with on-target amplicon.

### Off-target property of CRISPR-Cpf1 effector is addressed by CRISPR amplification

The Cpf1 effector targets thymine-rich regions in genes. Previous studies showed that CRISPR-Cpf1 induces less off-target mutations than CRISPR-Cas9 because it is less tolerant for mismatches^6, 7^. However, off-target mutations with rates below the NGS sensitivity limit (<0.5%) may occur, which can potentially cause serious problems, particularly in therapeutic settings. To address the safety issue, we assessed whether CRISPR amplification allows the detection of a small amount of Cpf1 induced off-target mutations with high sensitivity **(Fig. 2)**. A targeted sequence (*RPL32P3* site) in U2OS **(Fig. 2a, Supplementary Fig. 1b)** and HEK293FT cells **(Supplementary Fig. 2)** was analyzed for Cpf1-induced mutations measured as indel frequencies (0.1%∼11.2%) in off-target candidate loci predicted by *in-silico* analysis^18^ **(Supplementary Table 2)**. Ten predicted off-target sequences were processed using a optimally designed Cpf1-crRNA complex (ideally designed to cleave wild-type DNA for each target and off-target sites, respectively **(Supplementary Fig. 7a)**) to enrich the small amount of mutated DNA. The results indicated that mutant DNA amplification occurred at various indel frequency (99.9, 99.7, 97.5, 98.6 and 65.7% for off-target1-5, **Fig. 2a**), depending on the amount of off-target mutant DNA. The indel frequency for the on-target *RPL32P3* site **(Supplementary Table 2)** in U2OS cell was 7.7% **(Fig. 2a)**. After three rounds of CRISPR amplification, the mutant DNA fragment was amplified with an efficiency close to 100%(12.8 fold enriched). In a negative control treatment without Cpf1, the indel frequency could not be determined (∼0%) by three CRISPR amplifications. When off-target mutation detection by CRISPR amplification was compared with the conventional targeted amplicon sequencing^19^ or the result of GUIDE-seq technology^6^, candidate off-target sequences 1–5 **(Supplementary Table 2)** were commonly detected by CRISPR amplification (26.1, 41.2, 37, 155.5 and 117 fold enriched for off-target1-5), and the remaining candidate off-target sequences 6–10 **(Table 1)** did not show mutations **(Fig. 2b, Supplementary Table 4)**. In particular, CRISPR amplification was able to accurately identify off-target mutations (off-target loci 4(HEK293FT) and 5(U2OS)) that were difficult to identify by conventional targeted amplicon sequencing due to the low copy number of mutant DNA. The Cpf1-induced indel pattern revealed that off-target mutations were mainly deletions rather than insertions **(Supplementary Fig.4)**. In particular, amplification of insertions was not observed above 3^rd^-order enrichment. Overall, DNA fragments with large deletions around the cleavage site tended to be more effectively amplified than DNAs with smaller deletions. For CRISPR-Cpf1 off-target analysis at several loci, mutations were introduced in the *DNMT1* gene in HEK293FT cells **(Fig. 3a, Supplementary Fig. 7b)**. In addition to the indel frequency at the target sequence, the indel frequencies at two out of the three off-target genes predicted *in silico* **(Table 2)** were determined **(Fig. 3a, Supplementary Fig. 1c)**. In particular, indel mutations in off-target locus 2 that were difficult to detect by targeted amplicon sequencing were identified by CRISPR amplification **(Fig. 3a, b, Supplementary Table 5)**, and in off-target locus 1, no significant mutation was detected by this method. The indel pattern for each locus analyzed by sequencing revealed that deletions rather than insertions were the major enriched mutations in target and off-target sequences in the *DNMT1*-site3 locus **(Supplementary Fig. 5a)**.

**Fig. 2.**
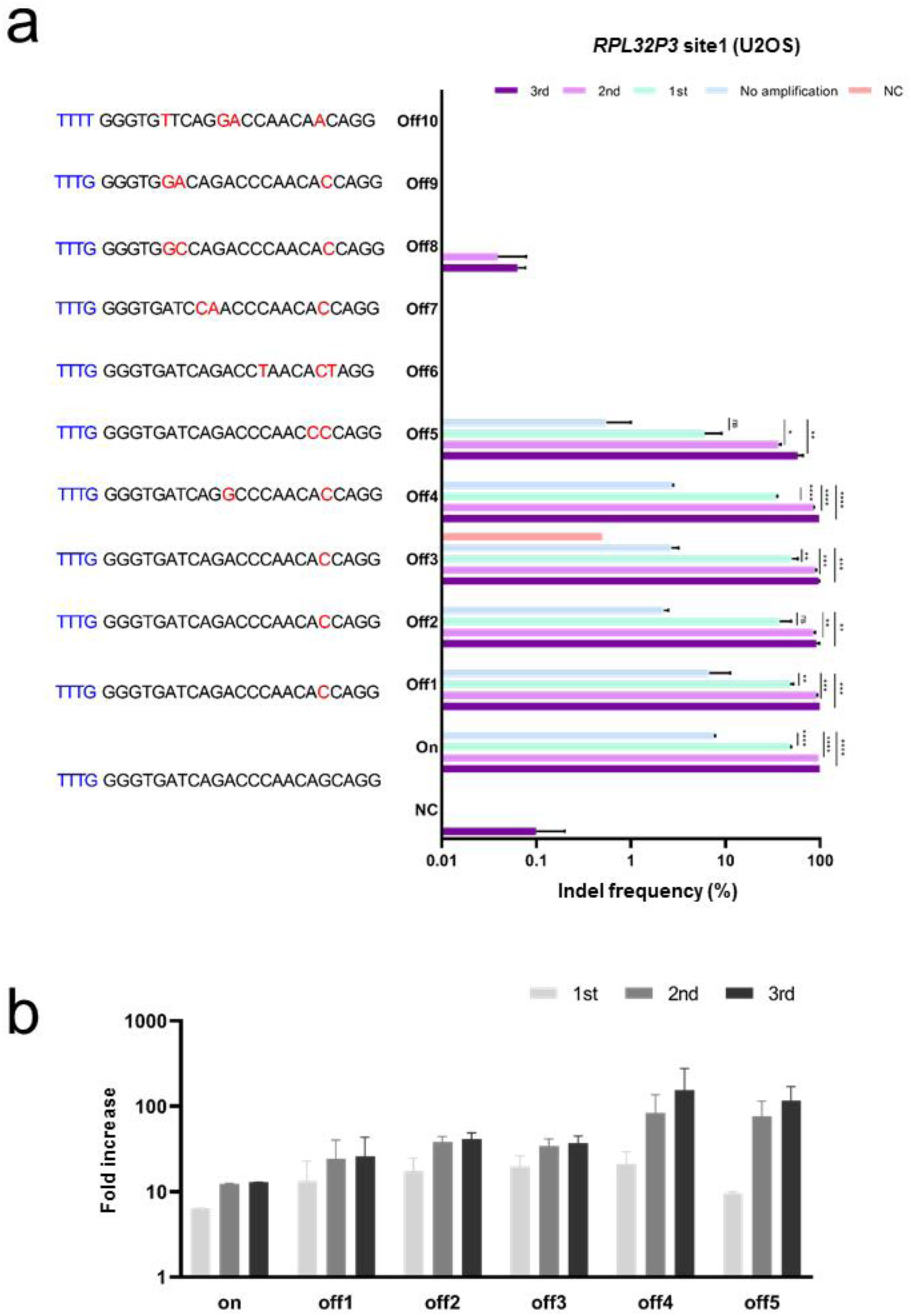
Detection of intracellular off-target mutations induced by CRISPR-Cpf1 by using CRISPR amplification. **(a)** Detection of off-target mutations for the target sequence (*RPL32P3* site) generated by the CRISPR-Cpf1 effector in U2OS cells. PCR amplicons were generated for 10 off-target sequences similar to the target sequence and the indel frequency (%) was determined by NGS after sequential CRISPR amplifications. Each round of CRISPR amplifications are marked by different colors. NC indicates a negative control for no Cpf1 delivery into the cells. The Y axis represents the amplified target and off-target sequences, and the X axis represents the indel frequency (%) in the analyzed amplicon on a log scale. *P*-values are calculated using a Tukey’s test(ns: not significant, *P*\*:<0.0332, *P*\*\*:<0.0021, *P*\*\*\*:<0.0002, *P*\*\*\*\*:<0.0001) **(b)** Fold increase after CRISPR amplification for each target and off-target sequence. Primary, secondary, and tertiary CRISPR amplification results are shown in gray, dark gray, and black, respectively. All experiments were conducted at least two times.

**Fig. 3.**
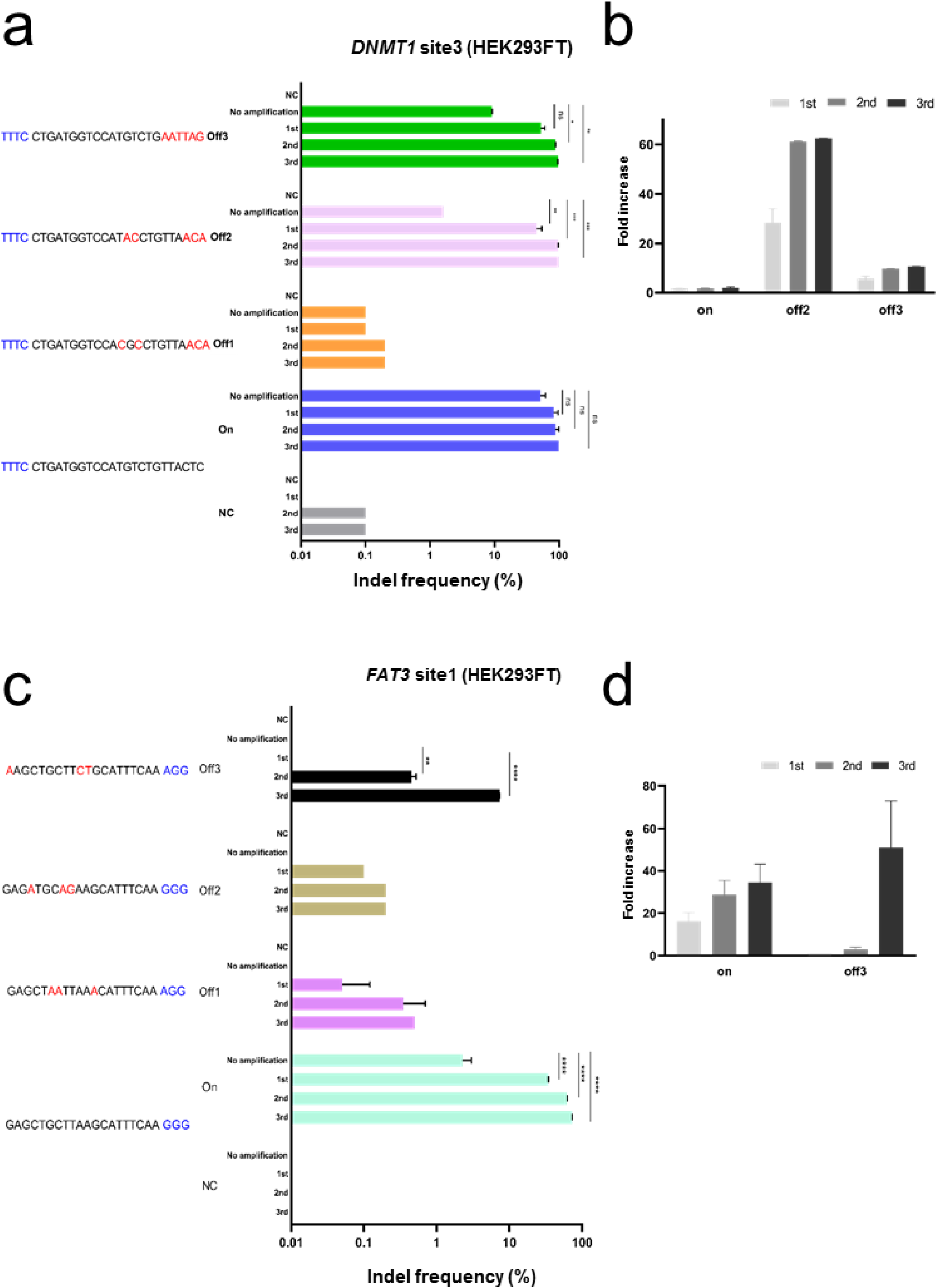
Detection of intracellular off-target mutations induced by CRISPR-Cpf1 and Cas9 by using CRISPR amplification. **(a)** Detection of off-target mutations for the target sequence (*DNMT1*) generated by the CRISPR-Cpf1 effector in HEK293FT cells. PCR amplicons were generated for three off-target sequences predicted *in silico* and the indel frequency (%) was analyzed by NGS after sequential CRISPR amplifications. *P*-values are calculated using a Tukey’s test(ns: not significant, *P*\*:<0.0332, *P*\*\*:<0.0021, *P*\*\*\*:<0.0002, *P*\*\*\*\*:<0.0001). **(b)** Fold increases in *DNMT1* target and off-target mutant DNA after CRISPR amplification. **(c)** Detection of off-target mutations for the target sequence (*FAT3* site) generated by the CRISPR-Cas9 effector in HEK293FT cells. PCR amplicons were generated for three off-target sequences predicted *in silico* and the indel frequency (%) was analyzed by NGS after sequential CRISPR amplifications. The Y axis represents the amplified target and off-target sequences, and the X axis represents the indel frequency (%) in the analyzed amplicon on a log scale. All experiments were conducted at least two times. *P*-values are calculated using a Tukey’s test(ns: not significant, *P*\*:<0.0332, *P*\*\*:<0.0021, *P*\*\*\*:<0.0002, *P*\*\*\*\*:<0.0001). **(d)** Fold increases in *FAT3* target and off-target mutant DNA after CRISPR amplification. In (b) and (d), primary, secondary, and tertiary CRISPR amplification results are shown in gray, dark gray, and black, respectively.

### Off-target detection for the CRISPR-Cas9 effector by CRISPR amplification

To verify the potential of the Cas9 effector for inducing off-target mutations at the cellular level, CRISPR amplification was applied to detect candidate off-target mutations **(Supplementary Table 2)** predicted based on the target sequence in HEK293FT edited with CRISPR-Cas9 **(Fig. 3c)**. In order to selectively amplify the mutated over wild-type DNA, the seed region of CRISPR-Cas9 target sequence was designed to harbor the PAM sequence (TTTN) of Cpf1 used for wild-type DNA cleavage **(Supplementary Fig. 7c)**. CRISPR amplification revealed a significant increase in indels in the target DNA, which was confirmed by NGS **(Fig. 3c)** and DNA cleavage assays **(Supplementary Fig. 3a)**. The amplification of on-target intracellular indels induced by CRISPR-Cas9 increased (34.4 fold) with consecutive rounds of CRISPR amplification **(Fig. 3d)**. The indel pattern showed a gradual increase in >2-bp deletions caused by CRISPR-Cas9 **(Supplementary Fig. 6a)**. A small amount of insertion mutations of various sizes was detected, but no significant amplification was detected. CRISPR amplification was also performed for the predicted off-target loci **(Supplementary Table 2)**, and unique indel patterns were detected at off-target site 3 **(Fig. 3c, d, Supplementary Fig. 6b)**. Conventional NGS did not allow detection of the off-target indels with frequencies below the detection limit. However, when the amplicon was subjected to third rounds of CRISPR amplification, even 1-bp-deletions were detected at a significant rate (51-fold increase vs. without amplification) **(Fig. 3d, Supplementary Fig. 6b, Supplementary Table 6)**.

### Single-base substituted off-target detection for adenine base editor (ABE) by CRISPR amplification

In addition to the insertions/deletions caused by target DNA cleavage by the CRISPR effector, we evaluated whether the CRISPR amplification can be used to detect single-base changes caused by off-target effects of adenine base editor (ABE) **(Fig. 4)**. To this end, first, ABE and gRNA **(Supplementary Table 1)** expression vectors were transfected into HEK293FT cells to induce single base substitutions at target genomic loci. Next, the genomic DNA extracted from the cells was subjected to CRISPR amplification for candidate off-target sequences predicted based on the gRNA sequence for the target DNA (*PSMB2*) **(Fig. 4a)**. To specifically amplify base-changed over wild-type DNA, the PAM sequence (TTTN) recognized by Cpf1 was placed in a window where the base substitution is mainly generated in the target sequence **(Supplementary Fig. 7d)**. If the PAM sequence recognized by Cpf1 has a base substitution, wild-type DNA without base substitution can be removed. Three cycles of CRISPR amplification on samples treated with ABE (*PSMB2* site) **(Supplementary Table 2)** revealed that DNA fragments with a base substitution (A>G) in the ABE editing window were significantly amplified (75.6% substitution, 7.67 fold increase) **(Fig. 4b)**. The base substitution (A>G) was found to occur mainly at adenine sites in the 12–18-bp window within the ABE target sequence **(Fig. 4c)**. The amount of the base substitution gradually increased with increasing cycles of CRISPR amplification. Next, CRISPR amplification was applied to predicted off-target sites **(Supplementary Table 2)**. The A>G base substitution was significantly induced in each of the off-target sequences (off-targets 2, 3, and 4 in **Fig. 4a, c**). A low amount of base substitution could not be detected by NGS, but it was detected by CRISPR amplification (434, 272.5 and 58 fold increase respectively) **(Fig. 4b, Supplementary Table 7)**. Interestingly, in contrast to target sequences, where the A>G base substitution occurred evenly within the window of the guide sequence (12–18 bp), intensive base substitutions occurred at the 18th adenine from the PAM sequence in off-target sequences as indicated by CRISPR amplification **(Fig. 4c)**.

**Fig. 4.**
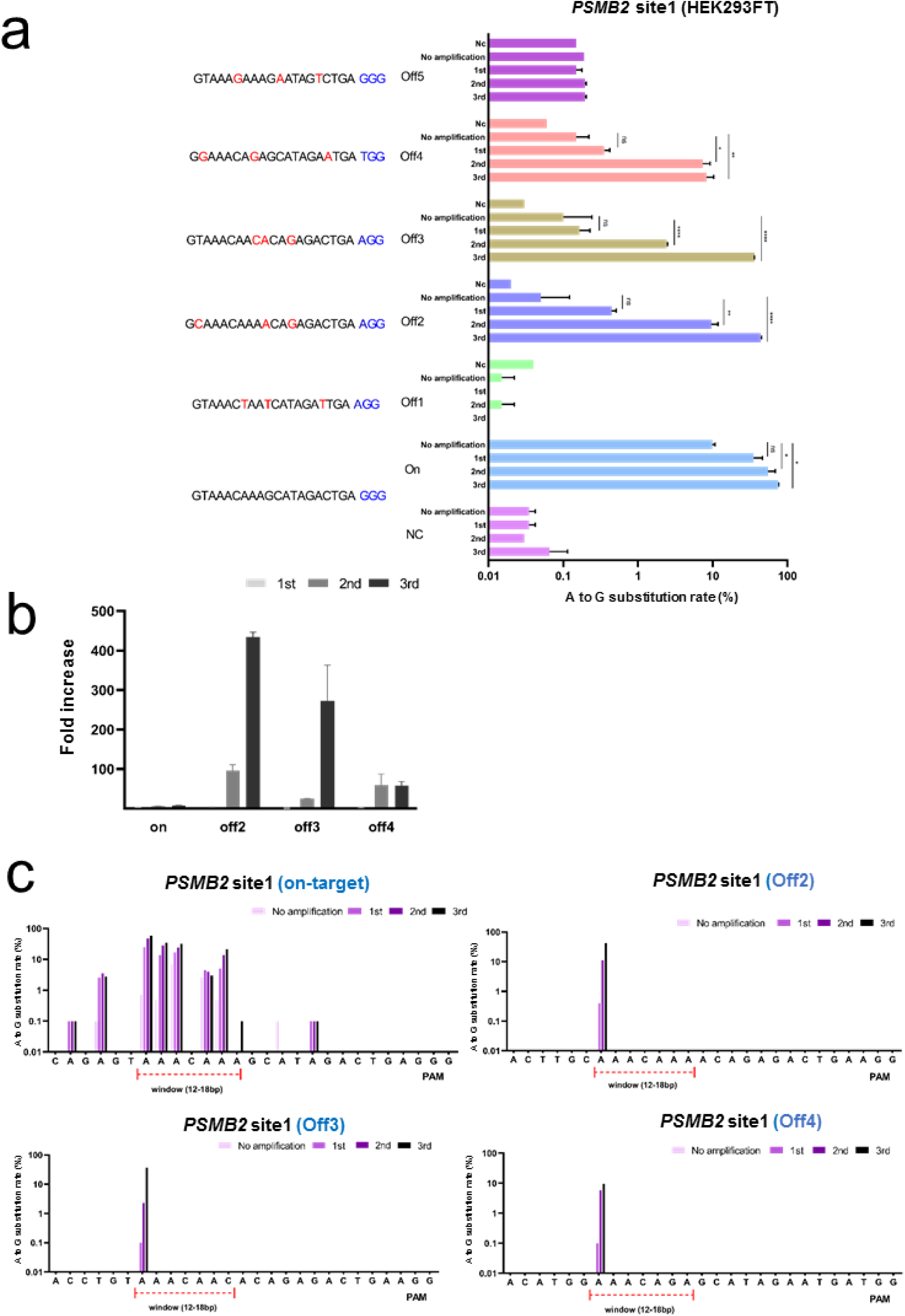
Detection of intracellular off-target mutations induced by ABE by using CRISPR amplification. (a) Detection of off-target mutations for the target sequence (*PSMB2* site) generated by ABE in HEK293FT cells. NGS was used to confirm whether single base mutations in target and non-target sequences can be amplified by CRISPR amplification. The Y axis represents the amplified target and off-target sequences, and the X axis represents the base substitution (A>G) frequency (%) in the analyzed amplicons on a log scale. All experiments were conducted at least two times. *P*-values are calculated using a Tukey’s test(ns: not significant, *P*\*:<0.0332, *P*\*\*:<0.0021, *P*\*\*\*:<0.0002, *P*\*\*\*\*:<0.0001). (b) Fold increases in single base substitution rates (%) after CRISPR amplification for target and off-target sequences. Primary, secondary, and tertiary CRISPR amplification results are shown in gray, dark gray, and black respectively. (c) Percentages of single base substitution frequency(%) in target and off-target sequences according to no, 1st, 2nd, and 3rd amplification, respectively. The target window is indicated in each panel.

## Discussion

As the CRISPR system is based on gRNAs that bind complementarily to target genes, it often causes mutations at unintended genomic loci with similar sequences. Efforts have been made to identify the presence of the off-target mutations, and to prevent their formation. In order to overcome the limitations regarding the sensitivity of current methods based on genome-wide analysis, we developed a method to amplify and detect small amounts of off-target mutations in intracellular genes generated by CRISPR effectors. The key of the mutated gene amplification method using CRISPR, developed in this study, is to enrich mutant genes containing off-target sequences predicted *in silico* and to remove the background wild-type DNA before PCR amplification.

As the CRISPR effectors, currently in use, sensitively recognize PAM sequences in the target DNA and the seed sequence of the protospacer, we considered these sequences when designing the gRNA. In particular, in the case of base editors that induce single-nucleotide substitutions, fragments containing mutations can be efficiently amplified by constructing a gRNA such that the PAM sequence is disrupted when a mutation is introduced. Our detection method using CRISPR amplification has various advantages. First, a very small amount of mutant DNA can be enriched in one round of CRISPR amplification, and stepwise amplification shows high sensitivity in detecting mutant DNA fragments of up to (0.00001%). Second, mutant genes can be easily amplified in a high-throughput manner. Third, as a gRNA and CRISPR effector are used, the method can be widely applied to insertions/deletions and single base substitutions generated by various CRISPR effectors. In addition, it is possible to apply the method to precisely analyze whether mutations are induced in off-target sequences by using new techniques, such as prime editing. One prerequisite of the current CRISPR amplification method is the need for accurate *in-silico* prediction of potential off-target sites, for which several online tools are available. Another technical difficulty is that there is a possibility of error incorporation during sequential PCR amplifications. In order to overcome this problem, Phusion High-Fidelity DNA Polymerase^20^ with an extremely low error rate was used, and a maximum of three amplification cycles was used based on a negative control experiment which does not shows significant enrichment of mutation patterns **(Fig. 2a, Fig. 3a,c, Fig. 4a)**.

Using the method developed in this study, we analyzed the off-target mutation propensity of CRISPR-Cpf1, CRISPR-Cas9, and ABE. The Cpf1 and Cas9 effectors respectively recognize T- and G-rich PAM sequences and bind to the sequences complementary to the gRNA in the target DNA to induce double-helix cleavage. These effectors are known to be very sensitive to mismatches in the PAM proximal region and less sensitive to mismatches in PAM distal regions. In this study, we compared CRISPR amplification with conventional NGS (<u>n</u>ext <u>g</u>eneration <u>s</u>equencing) or previously reported GUIDE-seq data for the detection of off-target mutations induced by CRISPR-Cpf1. Both methods revealed five off-target mutations for *RPL32P3* site in U2OS or HEK293FT cell, including various mutations in the PAM distal region. Our method allowed the amplification and detection of mutations present at levels below the NGS detection level. In addition, three off-target mutations in *DNMT1* were tested by our method. In particular, off-target indels with a very low read number in conventional NGS were effectively amplified by our CRISPR amplification method. Similar to the findings for CRISPR-Cpf1, when intracellular target mutation was induced by CRISPR-Cas9, mutations below the NGS detection level were detected at a high significance level in one of the *in-silico* predicted off-target loci. Surprisingly, in addition to the insertion/deletions caused by double-helix cleavage by CRISPR-Cpf1 or -Cas9, single-base substitutions at off-target sites induced by ABE were detected at very high significance levels. Using adequately designed gRNAs and CRISPR amplification, extremely small amounts of off-target base substitution (A>G) below the NGS detection level could be detected, and the sensitivity was augmented with increasing CRISPR amplification cycles.

The highly sensitive off-target mutation discovery method developed in this study can be combined with genome-wide methods for highly probable off-target candidate selection. After detecting the off-target candidates, CRISPR amplification can be applied to determine the authenticity of off-target candidates and to finally determine whether a specific gRNA can be used for gene therapy. In addition, it is possible to accurately analyze the tendency of existing gene editors to induce mutations that cannot be detected by NGS technologies and intracellular mutation patterns generated by newly developed gene editors. Finally, the CRISPR amplification technology can be applied to remove background wild-type DNA and enrich DNA mutations induced by newly developed gene editors, including prime editing, which will contribute to ultra-precision genome correction.

## Methods

### Protein purification

pET28-Cas12a (Cpf1) bacterial expression vectors were transformed into *Escherichia coli* BL21 (DE3) cells for the purification of Cpf1 recombinant protein (*Acidaminococcus sp.* (As)Cpf1). The cells were cultured at 37°C until they reached an optical density of 0.6. After 48 hours of isopropyl β-D-thiogalactoside induction, the cultures were centrifuged to remove the medium, and the cells were resuspended in buffer A (20 mM Tris-HCl (pH 8.0), 300 mM NaCl, 10 mM β-mercaptoethanol, 1% Triton X-100, 1 mM phenylmethylsulfonyl fluoride). Then, the cells were disrupted by sonication on ice for 3 min, and cell lysates were harvested by centrifugation at 5,000 rpm for 10 min. Ni-NTA resin was washed with buffer B (20 mM Tris-HCl (pH 8.0), 300 nM NaCl) and mixed with the cell lysates, and the mixtures were stirred for 1,5 h in a cold room (4°C). After centrifugation, non-specific binding components were removed by washing with 10 volumes of buffer B (20 mM Tris-HCl (pH 8.0), 300 nM NaCl), and buffer C (20 mM Tris-HCl (pH 8.0), 300 nM NaCl, 200 mM imidazole) was used to elute the Cpf1 protein bound to the Ni-NTA resin. The buffer was exchanged for buffer E (200 mM NaCl, 50 mM HEPES (pH 7.5), 1 mM DTT, 40% glycerol) using Centricon filters (Amicon Ultra) and the samples were aliquoted and stored at –80°C. Purity of the purified protein was confirmed by SDS-PAGE (10%), and protein activity was confirmed by an *in-vitro* PCR amplicon cleavage assay.

### *In-vitro* gRNA synthesis

For *in-vitro* transcription, DNA oligos containing a crRNA sequence **(Supplementary Table 1)** corresponding to each target sequence were purchased from Macrogen. crRNA was synthetized by incubating annealed (denaturation at 98°C for 30 s, primer annealing at 23°C for 30 s) template DNA was mixed with T7 RNA polymerase(NEB), 50 mM MgCl_2_, 100 mM rNTP (rATP, rGTP, rUTP, rCTP), 10× RNA polymerase reaction buffer, murine RNase inhibitor, 100 mM DTT, and DEPC at 37°C for 8 h. Thereafter, the DNA template was completely removed by incubation with DNase at 37°C for 1 h, and the RNA was purified using a GENECLEAN^®^ Turbo Kit (MP Biomedicals). The purified RNA was concentrated through lyophilization (2,000 rpm, 1 h, –55°C, 25°C).

### Cell culture and transfection

HEK293FT and U2OS human cells were obtained from the American Type Culture Collection. The cells were maintained in Dulbecco’s modified Eagle’s medium (DMEM) with 10% FBS (both from Gibco) at 37°C in the presence of 5% CO_2_. Cells were subcultured every 48 h to maintain 70% confluency. For target sequence editing, 10^5^ HEK293FT or U2OS cells were transfected with plasmids expressing crRNA (240 pmol) and Cpf1 (60 pmol) via electroporation using an Amaxa electroporation kit (V4XC-2032; program: CM-130 for HEK293FT, DN-100 for U2OS). Transfected cells were transferred to a 24-well plate containing DMEM (500 μl/well), pre-incubated at 37°C in the presence of 5% CO_2_ for 30 min, and incubated under the same conditions for subculture. After 48 h, genomic DNA was isolated using a DNeasy Blood & Tissue Kit (Qiagen).

### Genotyping by *in-vitro* cleavage

PCR amplicons were obtained from genomic DNA (from HEK293FT or U2OS) using DNA primers **(Supplementary Table 3)** corresponding to each target and non-target candidate gene locus. Purified recombinant Cpf1 and crRNA designed to remove DNA other than intracellularly induced mutant DNA were purified and premixed, and incubated in cleavage buffer (NEB3, 10 μl) at 37°C for 1 h. The reaction was stopped by adding a stop buffer (100 mM EDTA, 1.2% SDS), and DNA cleavage was confirmed by 2% agarose gel electrophoresis. DNA cleavage efficiency was calculated based on the cleaved band image pattern according to the formula: intensity of the cleaved fragment/total sum of the fragment intensity × 100%, using ImageJ software.

### Enrichment and validation of CRISPR-Cpf1 off-target mutations

A crRNA corresponding to the nucleotide sequence within the target gene was prepared **(Supplementary Table 1)**, and cells were transfected with plasmids expressing CRISPR-Cpf1 and the crRNA to induce genome mutation at the desired site. The target site (RPL32P3 site in human genomic DNA **(Supplementary Table 2)**) was selected considering the editing efficiency of Cpf1 at the given genome location. Cas-OFFinder^18^ was used to identify genome-wide candidate off-target sites derived from the target sequence **(Supplementary Table 2)**. To confirm the induction of off-target mutations by Cpf1, genomic DNA was extracted from cells incubated in the presence of plasmids expressing crRNA and Cpf1 for 48 h. Intracellular on-target and related candidate off-target genome sites were PCR-amplified (denaturation at 98°C for 30 s, primer annealing at 58°C for 30 s, elongation at 72°C for 30 sec, 35 cycles **(Supplementary Table 3)**). Non-mutated DNA fragments were removed by mixing crRNA **(Supplementary Table 1)**, Cpf1, and the amplicons, and incubating the mixture in cleavage buffer (NEB3, 10 μl) at 37°C for 1 h for complete cleavage of wild-type DNA. After this reaction, mutated DNA (targeted by intracellular treatment of Cpf1 and not cut by Cpf1 effector) was enriched by PCR using nested PCR primers, using only 2% (v/v) of the total reaction mixture. DNA fragments containing mutations (on- and off-target indels) are preferentially amplified over normal DNA. Amplification products were digested and analyzed by 2% agarose gel electrophoresis or NGS with barcode addition using nested PCR (denaturation at 98°C for 30 s, primer annealing at 58°C for 30 s, elongation at 72°C for 30 s, 35 cycles **(Supplementary Table 3)**).

### Enrichment and validation of CRISPR-Cas9 off-target mutations

To use CRISPR-Cas9 to induce mutations, a gRNA was designed and a gRNA expression plasmid was transfected into cells. The target was selected based on a high editing efficiency of the CRISPR-Cas9 effector (*FAT3* site in human genomic DNA **(Supplementary Table 2)**). Cas-OFFinder^18^ was used to identify genome-wide candidate off-target sites **(Supplementary Table 2)**. To confirm off-target mutation by CRISPR-Cas9, genomic DNA was extracted from transfected cells at 48 h after transfection. Intracellular on-target and related off-target candidate genomes site were PCR-amplified (denaturation at 98°C for 30 s, primer annealing at 58°C for 30 s, elongation at 72°C for 30 s, 35 cycles **(Supplementary Table 3)**). CRISPR amplification was performed using a crRNA (Table) designed to remove DNA other than mutant DNA. Subsequent steps were the same as described for CRISPR-Cpf1 off-target mutation analysis.

### Enrichment and validation of ABE off-target mutations

To use CRISPR-Cas9-based ABE to induce mutations, a gRNA complementary to the target sequence was prepared **(Supplementary Table 1).** gRNA and ABE expression vectors were simultaneously delivered into cells. The target sequence was selected based on a high editing efficiency of the ABE, and Cas-OFFinder^18^ was used to identify genome-wide candidate off-target sites **(Supplementary Table 2)**. Genomic DNA was extracted from the cells 96 h after transfection. Intracellular on-target and related off-target candidate genomes site were detected by PCR amplification (denaturation at 98°C for 30 s, primer annealing at 58°C for 30 s, elongation at 72°C for 30 sec, 35 cycles **(Supplementary Table 3)**). CRISPR amplification was performed by designing a crRNA **(Supplementary Fig. 7d, Supplementary Table1)** to remove DNA other than mutant DNA. Subsequent steps were the same as described for CRISPR-Cpf1 off-target mutation analysis.

### Targeted deep sequencing and data analysis

To confirm whether mutations were introduced in the target genome locus by the CRISPR-Cpf1, CRISPR-Cas9 or adenine base editor, genomic DNA extracted from cells was amplified using DNA primers **(Supplementary Table 3)** corresponding to target and off-target loci. Nested PCR (denaturation at 98°C for 30 s, primer annealing at 62°C for 15 s, elongation at 72°C for 15 s, 35 cycles) was performed to conjugate adapter and index sequences to the amplicons. Next, a barcoded amplicon mixture was loaded on mini-SEQ analyzer (Illumina MiniSeq system, SY-420-1001) and targeted deep sequencing was performed according to the manufacturer’s protocol. The Fastq data were analyzed with Cas-analyzer^21^, and the mutation frequency (%) was calculated(inserted and deleted allele frequency(%) / total allele frequency(%)).

## ACKNOWLEDGMENTS

This research was supported by a grant from NRF(2019R1C1C1006603) and KRIBB Research Initiative Program(KGM1051911, KGM4251824, KGM5381911).

