## Supplementary Table and Figures for "Prediction-based highly sensitive CRISPR off-target validation using target-specific DNA enrichment"

### Supplementary Tables

**Supplementary Table 1. Sequence information for crRNAs used in this study.** Direct repeat sequences of the AsCpf1 effector are shown in blue.

| target | crRNA Sequence | | |
|---|---|---|---|
| AGBL1 on | 5` | GAAUUUCUACUCUUGUAGAUGAUUGAAGGAAAGUUACAAAGG | 3` |
| RPL32P3 on | 5` | GAAUUUCUACUCUUGUAGAUGGGUGAUCAGACCCAACAGCAGG | 3` |
| RPL32P3 off1 | 5` | GAAUUUCUACUCUUGUAGAUGGGUGAUCAGACCCAACACCAGG | 3` |
| RPL32P3 off2 | 5` | GAAUUUCUACUCUUGUAGAUGGGUGAUCAGACCCAACACCAGG | 3` |
| RPL32P3 off3 | 5` | GAAUUUCUACUCUUGUAGAUGGGUGAUCAGACCCAACACCAGG | 3` |
| RPL32P3 off4 | 5` | GAAUUUCUACUCUUGUAGAUGGGUGAUCAGGCCCAACACCAGG | 3` |
| RPL32P3 off5 | 5` | GAAUUUCUACUCUUGUAGAUGGGUGAUCAGACCCAACCCCAGG | 3` |
| RPL32P3 off6 | 5` | GAAUUUCUACUCUUGUAGAUGGGUGAUCAGACCUAACACUAGG | 3` |
| RPL32P3 off7 | 5` | GAAUUUCUACUCUUGUAGAUGGGUGAUCCAACCCAACACCAGG | 3` |
| RPL32P3 off8 | 5` | GAAUUUCUACUCUUGUAGAUGGGUGGCCAGACCCAACACCAGG | 3` |
| RPL32P3 off9 | 5` | GAAUUUCUACUCUUGUAGAUGGGUGGACAGACCCAACACCAGG | 3` |
| RPL32P3 off10 | 5` | GAAUUUCUACUCUUGUAGAUGGGUGUUCAGGACCAACAACAGG | 3` |
| PSMB2 on | 5` | GAAUUUCUACUCUUGUAGAUCUCUGAGUGUACAAAAGAUGGUG | 3` |
| PSMB2 off2 | 5` | GAAUUUCUACUCUUGUAGAUCAAGUAUAUACCCAGUGCUGAGC | 3` |
| PSMB2 off3 | 5` | GAAUUUCUACUCUUGUAGAUCAGGUCACUAAAAAAUUAAAUGA | 3` |
| PSMB2 off4 | 5` | GAAUUUCUACUCUUGUAGAUCAUGUUUACACAUUAUUUAACUC | 3` |
| FAT3 on | 5` | GAAUUUCUACUCUUGUAGAUAAGGGAAGAAACCCUGAAACUCU | 3` |
| FAT3 off3 | 5` | GAAUUUCUACUCUUGUAGAUAAGGGGACAGCCACUACUGAGGC | 3` |
| DNMT1 site3 on | 5` | GAAUUUCUACUCUUGUAGAUCUGAUGGUCCAUGUCUGUUACUC | 3` |
| DNMT1 site3 off1 | 5` | GAAUUUCUACUCUUGUAGAUCUGAUGGUCCACGCCUGUUAACA | 3` |
| DNMT1 site3 off2 | 5` | GAAUUUCUACUCUUGUAGAUCUGAUGGUCCAUACCUGUUAACA | 3` |
| DNMT1 site3 off3 | 5` | GAAUUUCUACUCUUGUAGAUCUGAUGGUCCAUGUCUGAAUUAG | 3` |

**Supplementary Table 2. Sequence information for DNA targets amplified in this study.** PAM sequences (TTTN for AsCpf1 and NGG for SpCas9 effector) are shown in blue color and mismatched sequence to each on-target sites are shown in red color respectively.

| Target name | Mismatch | Sequence | Position | | Direction |
|---|---|---|---|---|---|
| AGBL1 on | - | **TTTA** GATTGAAGGAAAAGTTACAAAGG | chr15 | 86124495 | - |
| RPL32P3 on | - | **TTTG** GGGTGATCAGACCCAACAGCAGG | chr3 | 129389649 | - |
| RPL32P3 off1 | 1 | **TTTG** GGGTGATCAGACCCAACAcCAGG | chr8 | 108739813 | + |
| RPL32P3 off2 | 1 | **TTTG** GGGTGATCAGACCCAACAcCAGG | chr2 | 88944272 | - |
| RPL32P3 off3 | 1 | **TTTG** GGGTGATCAGACCCAACAcCAGG | chr6 | 33355560 | - |
| RPL32P3 off4 | 2 | **TTTG** GGGTGATCAGgCCCAACAcCAGG | chr1 | 205858150 | - |
| RPL32P3 off5 | 2 | **TTTG** GGGTGATCAGACCCAACccCAGG | chr15 | 78621826 | + |
| RPL32P3 off6 | 3 | **TTTG** GGGTGATCAGACCtAACActAGG | chr3 | 109693289 | - |
| RPL32P3 off7 | 3 | **TTTG** GGGTGATCcaACCCAACAcCAGG | chr4 | 43472802 | - |
| RPL32P3 off8 | 3 | **TTTG** GGGTGgcCAGACCCAACAcCAGG | chr19 | 53480825 | + |
| RPL32P3 off9 | 3 | **TTTG** GGGTGgaCAGACCCAACAcCAGG | chr20 | 42901255 | - |
| RPL32P3 off10 | 4 | **TTTT** GGGTGtTCAGgaCCAACAaCAGG | chr12 | 133092583 | + |
| PSMB2 on | - | GTAAACAAAGCATAGACTGA **GGG** | chr1 | 35631454 | + |
| PSMB2 off2 | 3 | GcAAACAAAaCAgAGACTGA **AGG** | chr3 | 77072677 | - |
| PSMB2 off3 | 3 | GTAAACAAcaCAgAGACTGA **AGG** | chr7 | 112493828 | - |
| PSMB2 off4 | 3 | GgAAACAgAGCATAGAaTGA **TGG** | chr1 | 208960762 | + |
| FAT3 on | - | GAGCTGCTTAAGCATTTCAA **GGG** | chr11 | 92532744 | - |
| FAT3 off3 | 3 | GAGaTGCagAAGCATTTCAA **GGG** | chr18 | 25854400 | + |
| DNMT1 site3 on | - | **TTTC** CTGATGGTCCATGTCTGTTACTC | chr19 | 10,133,766 | - |
| DNMT1 site3 off1 | 5 | **TTTC** CTGATGGTCCAcGcCTGTTAaca | chrX | 98291178 | - |
| DNMT1 site3 off2 | 5 | **TTTC** CTGATGGTCCATacCTGTTAaca | chrX | 93421364 | - |
| DNMT1 site3 off3 | 6 | **TTTC** CTGATGGTCCATGTCTGaattag | chr1 | 213204013 | + |

**Supplementary Table 3. Sequence information for DNA primers used in this study.** Primer sequence information to amplify the target gene was indicated. The base sequences of the forward and reverse adapter primers used in the next generation sequencing are shown in green and blue, respectively.

| Target gene (primer direction) | DNA sequence (5' to 3') |
|---|---|
| AGBL1 on-target F1 | TGCCCTGCCATTATATAGACTGTT |
| AGBL1 on-target R1 | AAAATACAAAATGTAGCGGGGCA |
| AGBL1 on-target F2 | CATTACAACCTTCTTCTGTTTGTC |
| AGBL1 on-target R2 | ACCAGAGTGAGACTCTGTCT |
| FAT3 on-target F1 | AGGAGTTTGAAGCTGTAGTAAG |
| FAT3 on-target F2 | CGGGAGACTCTGTCTCTTAA |
| FAT3 on-target R1 | AACCCGCCTTTTGTTACAAGTT |
| FAT3 on-target R2 | CAAAACTTCAAACAGAATACATGTG |
| FAT3 off-target3 F1 | GAGAATACTGTGCAGCCAGAG |
| FAT3 off-target3 F2 | AGAAGTTGAGAGACCCTCAGAA |
| FAT3 off-target3 R1 | CCAGCACACCTTGGTAATCTG |
| FAT3 off-target3 R2 | GTAACCTCTCCTAACCAGCACA |
| RPL32P3 on-target F1 | ACCATGTCGTCCTGTATGATC |
| RPL32P3 on-target F2 | CTTCCTGAAGACACCGATTCT |
| RPL32P3 on-target R1 | AGAGACCAACAATGACCTGTTT |
| RPL32P3 on-target R2 | AGAACCTTGCACATGAAATGTG |
| RPL32P3 off-target1 F1 | CAGCAATTGCTATCTGTTGTAGTT |
| RPL32P3 off-target1 F2 | TCTGTTGTAGTTCTTGTGGGTT |
| RPL32P3 off-target1 R1 | AGTGTTGGGAAGAAAGCTGAG |
| RPL32P3 off-target1 R2 | TCCTCTGGAATGTGTCTAGACT |
| RPL32P3 off-target2 F1 | AGTCTCCATTCTGTTCATGCC |
| RPL32P3 off-target2 F2 | TTCATGCCCACAATGGTGATAT |
| RPL32P3 off-target2 R1 | CTCTCCAAGGATCGATTGTATCT |
| RPL32P3 off-target2 R2 | CTGCTCTCCATTATCTCAAGTA |
| RPL32P3 off-target3 F1 | GTAAGGGTGCACCCTTCTATA |
| RPL32P3 off-target3 F2 | GTACCTAGCCTTGCTGAGAA |
| RPL32P3 off-target3 R1 | CCTTTGGAATGTGTCCAGAC |
| RPL32P3 off-target3 R2 | TGCTGGCTTCTTGCTTCTAG |
| RPL32P3 off-target4 F1 | GCAAAACTGCTCCACTGTACT |
| RPL32P3 off-target4 F2 | ACCTTGGTTCTTCAGAGTGC |
| RPL32P3 off-target4 R1 | TGTTGGTTCCTTGCTTCTTACTT |
| RPL32P3 off-target4 R2 | CCTCCATTATCTCAAGCAGCA |
| RPL32P3 off-target5 F1 | GTGACAGAGTGAGACTTCATCT |
| RPL32P3 off-target5 F2 | GGTGGGTGGAAGTTAGTTGA |
| RPL32P3 off-target5 R1 | ACATTCTTGGGGTTGCAGATG |
| RPL32P3 off-target5 R2 | CTCTCAGATTGCTCCATCAGAA |
| RPL32P3 off-target6 F1 | GTAGAGAACTGAAGAAAGATCTGC |
| RPL32P3 off-target6 F2 | AGAGCAGAGGTCTCCATTCTTA |
| RPL32P3 off-target6 R1 | CACTATTATTGCGTCTAAGTCCTT |
| RPL32P3 off-target6 R2 | CTGGGGTCTAAGGTCTAAAAC |
| RPL32P3 off-target7 F1 | CTATGGGATACAGCCAAAACAGT |
| RPL32P3 off-target7 F2 | ACAGCCAAAACAGTACTAAGAG |
| RPL32P3 off-target7 R1 | TGATAGCTGTGGTGGTTTTACAA |
| RPL32P3 off-target7 R2 | AGCTGAGTGTTGGGAGAGAA |
| RPL32P3 off-target8 F1 | TCAGTGCCTACCTGGTAAAGAT |
| RPL32P3 off-target8 F2 | GTCATCTGTGGACATCTTGAG |
| RPL32P3 off-target8 R1 | TGATCTCAAGTCGCAGAACAT |
| RPL32P3 off-target8 R2 | AAGTCGCAGAACATGTTCCATA |
| RPL32P3 off-target9 F1 | GTGGACATACACAAGTGTTTCA |
| RPL32P3 off-target9 F2 | GTATGACATTCTGTGATGGGAG |
| RPL32P3 off-target9 R1 | CCAGGACTGATTGTATCTTGAGT |
| RPL32P3 off-target9 R2 | CACTATCTCAAGCAGCAGAACA |
| RPL32P3 off-target10 F1 | CTAACAGAATACGATGCTGCC |

| | |
|---|---|
| RPL32P3 off-target10 F2 | TACGATGCTGCCAATCAAAAGT |
| RPL32P3 off-target10 R1 | GCTGATACTGCAGTGGTACT |
| RPL32P3 off-target10 R2 | TTCACAGGCTCCATCTCTCT |
| PSMB2 on-target F1 | CTTCATATTGGTGTGTCCCAAC |
| PSMB2 on-target F2 | AATGAACAAGTAGCAACAGGAGG |
| PSMB2 on-target R1 | AGCTGGGATTACAGGCATGTAC |
| PSMB2 on-target R2 | TCCCAAAGCTCTAGGATTACAG |
| PSMB2 off-target2 F1 | GCCTAGCAGATTTATTTTCTGTTC |
| PSMB2 off-target2 F2 | TTCTGACTTCTGCTACTCATTGC |
| PSMB2 off-target2 R1 | ACAAACTCAAGACTGTCACTGATT |
| PSMB2 off-target2 R2 | GGTGGGGAACTTGTGATCTAG |
| PSMB2 off-target3 F1 | ACCCGAGCAGCACTACTTTTC |
| PSMB2 off-target3 F2 | TGCTTGCTCAAACCTGCTATC |
| PSMB2 off-target3 R1 | CTACAAAGGGTGTCAGAGGCA |
| PSMB2 off-target3 R2 | GGTGGGACTAGGAATTCCTG |
| PSMB2 off-target4 F1 | CCAAAGGAAGATACAGCAGTGT |
| PSMB2 off-target4 F2 | GTAGTCAACCTCAGTGTCCAT |
| PSMB2 off-target4 R1 | AATGCATTTCTGGTTACCCTGTT |
| PSMB2 off-target4 R2 | GTGTGAAGTGGTCAACTACAAG |
| DNMT1-site3 on-target F1 | CAAAGCCATTGGCTTGGAGAT |
| DNMT1-site3 on-target F2 | AGATCAAGCTTTGTATGTTGGCC |
| DNMT1-site3 on-target R1 | AGAAGTCCCGTGCAAATCAC |
| DNMT1-site3 on-target R2 | GCAAATCACGAATACCCACCC |
| DNMT1-site3 off-target1 F1 | TGTTGCAAGTCCCATGAGGA |
| DNMT1-site3 off-target1 F2 | CAGGGAACTCTAATCTCACAAT |
| DNMT1-site3 off-target1 R1 | GAACAGGAAAGAAAGGAAAATGAG |
| DNMT1-site3 off-target1 R2 | CCTCTTTCCCATGATTCTTCC |
| DNMT1-site3 off-target2 F1 | AGCAGGTCATTGGCAATGATAC |
| DNMT1-site3 off-target2 F2 | CAAATGTTTGTGCAGGTTGATGTT |
| DNMT1-site3 off-target2 R1 | GGTTTAGAGCAGGAGTGAAAGT |
| DNMT1-site3 off-target2 R2 | AAGAAAGGAAAGTTCACTTGGAAG |
| DNMT1-site3 off-target3 F1 | AACCCTGCTACCTACTGAGAAT |
| DNMT1-site3 off-target3 F2 | CATTAGCCTGTGTTTTCACATAAG |
| DNMT1-site3 off-target3 R1 | TTAATAGCATCAAAGGCAAACCAT |
| DNMT1-site3 off-target3 R2 | ATGATGGGAAAGTGTGCAAATAG |
| AGBL1_on_Adaptor_F | ACACTCTTTCCCTACACGACGCTCTTCCGATCT CTTAAGGAAACAGAAGAGAAATCTGCGTG |
| AGBL1_on_Adaptor_R | GTGACTGGAGTTCAGACGTGTGCTCTTCCGATCT ATTATAAGATGTTGACAATAACACAACAGG |
| FAT3_on_Adaptor_F | ACACTCTTTCCCTACACGACGCTCTTCCGATCT GTGGATTGTTTCATTCCAATGATGAAACAA |
| FAT3_on_Adaptor_R | GTGACTGGAGTTCAGACGTGTGCTCTTCCGATCT CATTTTGAACAGCCTTCTCCAGAAAACA |
| FAT3_off3_Adaptor_F | ACACTCTTTCCCTACACGACGCTCTTCCGATCT GGAAAATGAGCACTTAGCAATCAATTTG |
| FAT3_off3_Adaptor_R | GTGACTGGAGTTCAGACGTGTGCTCTTCCGATCT CTGTTTCTTTTTCTTGTTACTGCAGCA |
| RPL32P3_on_Adaptor_F | ACACTCTTTCCCTACACGACGCTCTTCCGATCT TGGGACAGGGTGGGTTACCTTG |
| RPL32P3_on_Adaptor_R | GTGACTGGAGTTCAGACGTGTGCTCTTCCGATCT ATCCTCACCACCTGTTTGTTGCA |
| RPL32P3_off1_Adaptor_F | ACACTCTTTCCCTACACGACGCTCTTCCGATCT TAGCTAGGGTGACTTGGCTAGCTTG |
| RPL32P3_off1_Adaptor_R | GTGACTGGAGTTCAGACGTGTGCTCTTCCGATCT GGTCCCACTTCTCTCTCTCAAATTGT |
| RPL32P3_off2_Adaptor_F | ACACTCTTTCCCTACACGACGCTCTTCCGATCT AAGCCCTGCAAATGTAAAAATCATAACA |
| RPL32P3_off2_Adaptor_R | GTGACTGGAGTTCAGACGTGTGCTCTTCCGATCT TGCCCACATTTCTCTCTCAAACTGTC |
| RPL32P3_off3_Adaptor_F | ACACTCTTTCCCTACACGACGCTCTTCCGATCT AAAAATCAAGACCTACCCAGTGCAAG |
| RPL32P3_off3_Adaptor_R | GTGACTGGAGTTCAGACGTGTGCTCTTCCGATCT |

| | |
|---|---|
| | CTTTGCAACCTCCATACTAGTATTGGC |
| RPL32P3_off4_Adaptor_F | ACACTCTTTCCCTACACGACGCTCTTCCGATCT AGCATTTCTTTACCATGGTCTTCATAGC |
| RPL32P3_off4_Adaptor_R | GTGACTGGAGTTCAGACGTGTGCTCTTCCGATCT TTCGCAGCCTCCAATCTAGTGTT |
| RPL32P3_off5_Adaptor_F | ACACTCTTTCCCTACACGACGCTCTTCCGATCT GTACCTGGAGGTTTGTTATACTGTTCTCT |
| RPL32P3_off5_Adaptor_R | GTGACTGGAGTTCAGACGTGTGCTCTTCCGATCT GCCTGGTCCCGCTTTCTCTCTT |
| RPL32P3_off6_Adaptor_F | ACACTCTTTCCCTACACGACGCTCTTCCGATCT ATGCATCCATGTATATTTTGGAGCATTCA |
| RPL32P3_off6_Adaptor_R | GTGACTGGAGTTCAGACGTGTGCTCTTCCGATCT TCTTTTTTGCAAAGACCCACCTTATGT |
| RPL32P3_off7_Adaptor_F | ACACTCTTTCCCTACACGACGCTCTTCCGATCT ACAAAACTGAACAAACCTTTAGCCAGA |
| RPL32P3_off7_Adaptor_R | GTGACTGGAGTTCAGACGTGTGCTCTTCCGATCT GCAGCCTCCATACTAGCATTGG |
| RPL32P3_off8_Adaptor_F | ACACTCTTTCCCTACACGACGCTCTTCCGATCT GCTATTAAGTGCATGTTTCCCTCAAGG |
| RPL32P3_off8_Adaptor_R | GTGACTGGAGTTCAGACGTGTGCTCTTCCGATCT CTGGACCCACCTTATGCATTCTTAACT |
| RPL32P3_off9_Adaptor_F | ACACTCTTTCCCTACACGACGCTCTTCCGATCT AGGCACTGTGCTGTCTTACATGTTATT |
| RPL32P3_off9_Adaptor_R | GTGACTGGAGTTCAGACGTGTGCTCTTCCGATCT CCTGGACACACTTTCTCTCTCAAACT |
| RPL32P3_off10_Adaptor_F | ACACTCTTTCCCTACACGACGCTCTTCCGATCT TATGGCTTGTGATTTCTTGTTTCCATAACT |
| RPL32P3_off10_Adaptor_R | GTGACTGGAGTTCAGACGTGTGCTCTTCCGATCT AGACCAGGTTTACAGAAGGCTAGAA |
| PSMB2_on_Adaptor_F | ACACTCTTTCCCTACACGACGCTCTTCCGATCT AGTAAAGCAGAAGGAATAACAGTGCCC |
| PSMB2_on_Adaptor_R | GTGACTGGAGTTCAGACGTGTGCTCTTCCGATCT TTCTGTTTGTGGCCAAGAATTGCTGT |
| PSMB2_off2_Adaptor_F | ACACTCTTTCCCTACACGACGCTCTTCCGATCT ACACAAACCAAGGGTGATGAAGTTT |
| PSMB2_off2_Adaptor_R | GTGACTGGAGTTCAGACGTGTGCTCTTCCGATCT CATAATTTCGAAACATATACACAATGGG |
| PSMB2_off3_Adaptor_F | ACACTCTTTCCCTACACGACGCTCTTCCGATCT GACAAGAGATTACTAGTGTTGCCTAAACA |
| PSMB2_off3_Adaptor_R | GTGACTGGAGTTCAGACGTGTGCTCTTCCGATCT TCAGCATTTTCTTCTGTATCATGGGAG |
| PSMB2_off4_Adaptor_F | ACACTCTTTCCCTACACGACGCTCTTCCGATCT GGAGGACATTATCTTAAATGAAACAACTC |
| PSMB2_off4_Adaptor_R | GTGACTGGAGTTCAGACGTGTGCTCTTCCGATCT CCACATTGAACCCAACAAGCAACTT |
| DNMT1-site3_on_Adaptor_F | ACACTCTTTCCCTACACGACGCTCTTCCGATCT GTTGCACGTGTCAAGTGCTTAGAG |
| DNMT1-site3_on_Adaptor_R | GTGACTGGAGTTCAGACGTGTGCTCTTCCGATCT GACTGAACACTCCTCAAACGGTC |
| DNMT1-site3_off1_Adaptor_F | ACACTCTTTCCCTACACGACGCTCTTCCGATCT CTACCCCCACCACTAGAAATGCCA |
| DNMT1-site3_off1_Adaptor_R | GTGACTGGAGTTCAGACGTGTGCTCTTCCGATCT TTTGTCTCTTAACATGCATGCCTAGGAA |
| DNMT1-site3_off2_Adaptor_F | ACACTCTTTCCCTACACGACGCTCTTCCGATCT CCTCAACCACTAAATATGTTATTTAGTGGT |
| DNMT1-site3_off2_Adaptor_R | GTGACTGGAGTTCAGACGTGTGCTCTTCCGATCT CCTTTTCCGATGGAGTGTACTCAGAAGAT |
| DNMT1-site3_off3_Adaptor_F | ACACTCTTTCCCTACACGACGCTCTTCCGATCT GGTCTCAAATAAGTTTGAGAATGAATGTG |
| DNMT1-site3_off3_Adaptor_R | GTGACTGGAGTTCAGACGTGTGCTCTTCCGATCT AGAGCTGAAAGTTTAGCATGGAGG |

| Target sequence | GUIDE-seq[6] (Reported data) | Targeted amplicon sequencing (NGS, HEK293FT) | CRISPR enrichment (NGS, HEK293FT) | CRISPR enrichment (NGS, U2OS) |
|---|---|---|---|---|
| **RPL32P3-on target** (TTTGGGGTGATCAGACCCAACAGCAGG) | O | O | O | O |
| **RPL32P3-off target1** (TTTGGGGTGATCAGACCCAACAcCAGG) | O | O | O | O |
| **RPL32P3-off target2** (TTTGGGGTGATCAGACCCAACAcCAGG) | O | O | O | O |
| **RPL32P3-off target3** (TTTGGGGTGATCAGACCCAACAcCAGG) | O | O | O | O |
| **RPL32P3-off target4** (TTTGGGGTGATCAGgCCCAACAcCAGG) | O | O | O | O |
| **RPL32P3-off target5** (TTTGGGGTGATCAGACCCAACccCAGG) | O | ? | O | O |
| **RPL32P3-off target6** (TTTGGGGTGATCAGACCtAACActAGG) | X | X | X | X |
| **RPL32P3-off target7** (TTTGGGGTGATCcaACCCAACAcCAGG) | X | X | X | X |
| **RPL32P3-off target8** (TTTGGGGTGgcCAGACCCAACAcCAGG) | X | X | X | X |
| **RPL32P3-off target9** (TTTGGGGTGgaCAGACCCAA | X | X | X | X |

| | | | | |
|---|---|---|---|---|
| CAcCAGG) | | | | |
| **RPL32P3-off target10** (**TTTT**GGGTGtTCAGgaCCAACAaCAGG) | X | X | X | X |

**Supplementary Table 4. Comparative analysis of the CRISPR amplification method and a conventional method (GUIDE-seq) for the detection of intracellular off-target mutations induced by CRISPR-Cas12a (Cpf1).** PAM sequences(TTTN) of the AsCpf1 effector are shown in blue and mismatched sequence to each on-target sites are shown in red respectively. (O) indicates detection, (X) indicates no detection and (?) indicates ambiguous detection (indel frequency(%) below the detection limit <0.5%) with applied methods. NGS: finally confirmed by next generation sequencing. Yellow mark indicates the mutant DNA detection with CRISPR amplification methods not by conventional NGS methods.

| Target sequence | Targeted amplicon sequencing (NGS, HEK293FT) | CRISPR enrichment (NGS, HEK293FT) |
|---|---|---|
| **DNMT1-site3-on target** (**TTTC**CTGATGGTCCATGTCTGTTACTC) | O | O |
| **DNMT1-site3-off target1** (**TTTC**CTGATGGTCCAcGcCTGTTAaca) | X | X |
| **DNMT1-site3-off target2** (**TTTC**CTGATGGTCCATacCTGTTAaca) | O | O |
| **DNMT1-site3-off target3** (**TTTC**CTGATGGTCCATGTCTGaattag) | O | O |

**Supplementary Table 5. Comparative analysis of the CRISPR amplification method and a conventional method (targeted amplicon sequencing) for the detection of intracellular off-target mutations induced by CRISPR-Cpf1.** PAM sequences(TTTN) of the AsCpf1 effector are shown in blue and mismatched sequence to each on-target sites are shown in red respectively. (O) indicates detection and (X) indicates no detection with applied methods. NGS: finally confirmed by next generation sequencing.

| Target sequence | Targeted amplicon sequencing (NGS, HEK293FT) | CRISPR enrichment (NGS, HEK293FT) |
|---|---|---|
| **FAT3-on target** (GAGCTGCTTAAGCATTTCAA**GGG)** | O | O |
| **FAT3-off target2** (GcAAACAAAaCAgAGACTGA**AGG)** | X | X |
| **FAT3-off target3** (GTAAACAAcaCAgAGACTGA**AGG)** | X | O |
| **FAT3-off target4** (GgAAACAgAGCATAGAaTGA**TGG)** | X | X |

**Supplementary Table 6. Comparative analysis of the CRISPR amplification method and a conventional method (targeted amplicon sequencing) for the detection of intracellular off-target mutations induced by CRISPR-Cas9.** PAM sequences(NGG) of the SpCas9 effector are shown in blue and mismatched sequence to each on-target sites are shown in red respectively. (O) indicates detection and (X) indicates no detection with applied methods. NGS: finally confirmed by next generation sequencing.Yellow mark indicates the mutant DNA detection with CRISPR amplification methods not by conventional NGS methods.

| Target sequence | Targeted amplicon sequencing (NGS, HEK293FT) | CRISPR enrichment (NGS, HEK293FT) |
|---|---|---|
| **PSMB2-on target** (GTAAACAAAGCATAGACTGA**GGG**) | O | O |
| **PSMB2-off target2** (GcAAACAAAaCAgAGACTGA**AGG**) | X | O |
| **PSMB2-off target3** (GTAAACAAcaCAgAGACTGA**AGG**) | X | O |
| **PSMB2-off target4** (GgAAACAgAGCATAGAaTGA**TGG**) | X | O |

**Supplementary Table 7. Comparative analysis of the CRISPR amplification method and a conventional method (targeted amplicon sequencing) for the detection of intracellular off-target mutations induced by ABE.** PAM sequences(NGG) of the adenine base editor are shown in blue and mismatched sequence to each on-target sites are shown in red respectively. (O) indicates detection and (X) indicates no detection with applied methods. NGS: finally confirmed by next generation sequencing. Yellow mark indicates the mutant DNA detection with CRISPR amplification methods not by conventional NGS methods.

### Supplementary Figures

Supplementary Fig.1

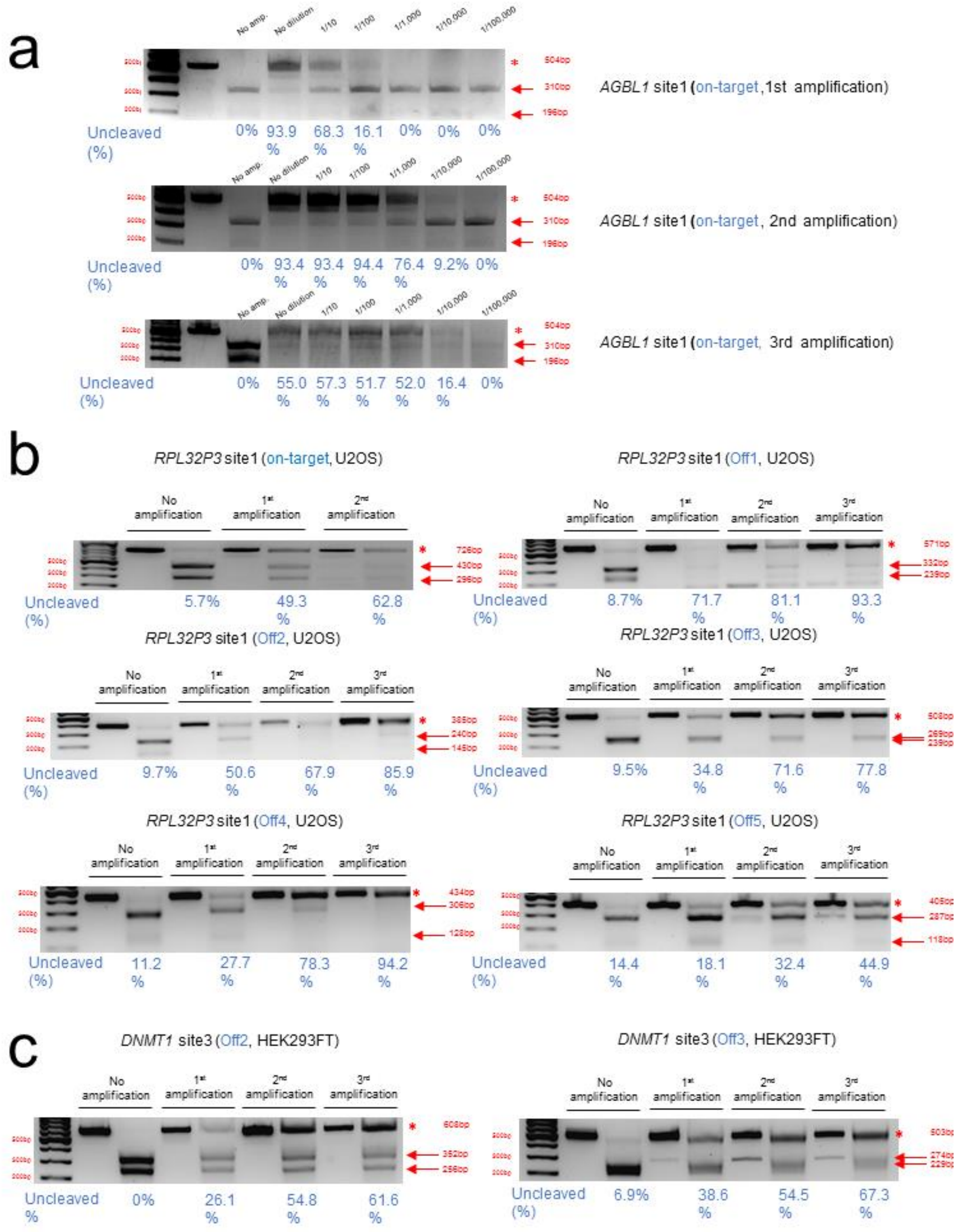


**Supplementary Figure 1. Genotyping of the Cpf1 induced mutant DNA enrichment with CRISPR amplification. (a)** Relatively compare the results of serially diluted (from 1X to 1/ 100000X) each genomic DNA sample with mutations induced by Cpf1. Relative mutant DNA frequency(%) at *AGBL1* locus was calculated by cleaving DNA amplicons with optimally designed crRNA for Cpf1. The cleaved amplicons were separated by 2% agarose gel. **(b)** Relative mutant DNA frequency(%) for predicted off-target sites of *RPL32P3* site in genomic DNA from U2OS cells. **(c)** Relative mutant DNA frequency(%) for predicted off-target sites of *DNMT1* in genomic DNA from U2OS cells. Uncleaved DNA fractions are indicated by asterisk and cleaved DNA fractions are indicated by red arrows.

Supplementary Fig.2

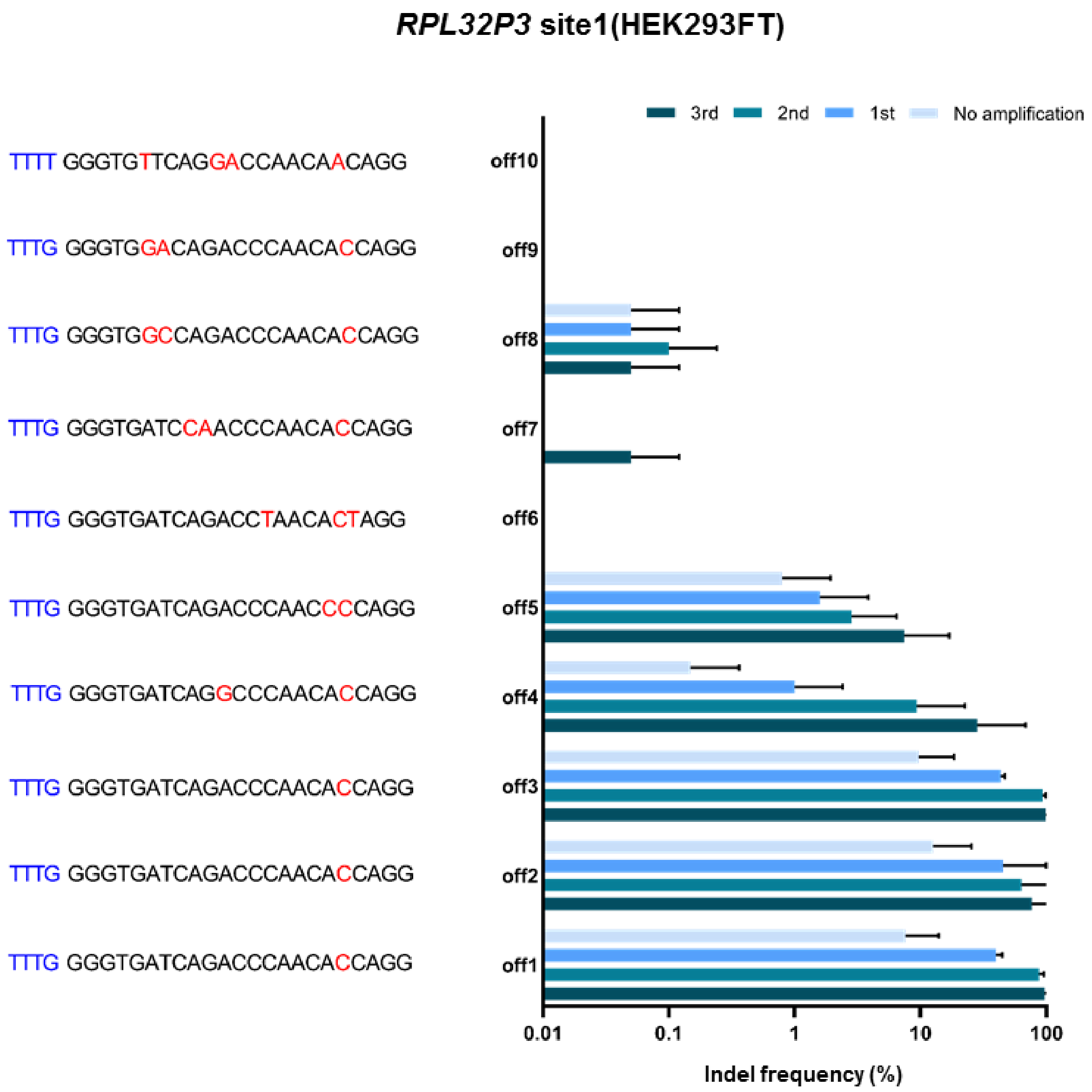


**Supplementary Figure 2. Detection of the intracellular off-target mutation induced by CRISPR-Cas12a(Cpf1) by using CRISPR amplification.** Detection of off-target mutations for the target sequence (*RPL32P3* locus) generated by CRISPR-Cpf1 effector in HEK293FT cells. PCR amplicons were generated for 10 off-target sequences same with **(Fig. 2)** and indel frequency(%) was analyzed by next-generation sequencing after multiple CRISPR amplification. The Y axis represents the amplified target and off-target sequences, and the X axis represents the frequency (%) of indels on a log scale.

Supplementary Fig.3

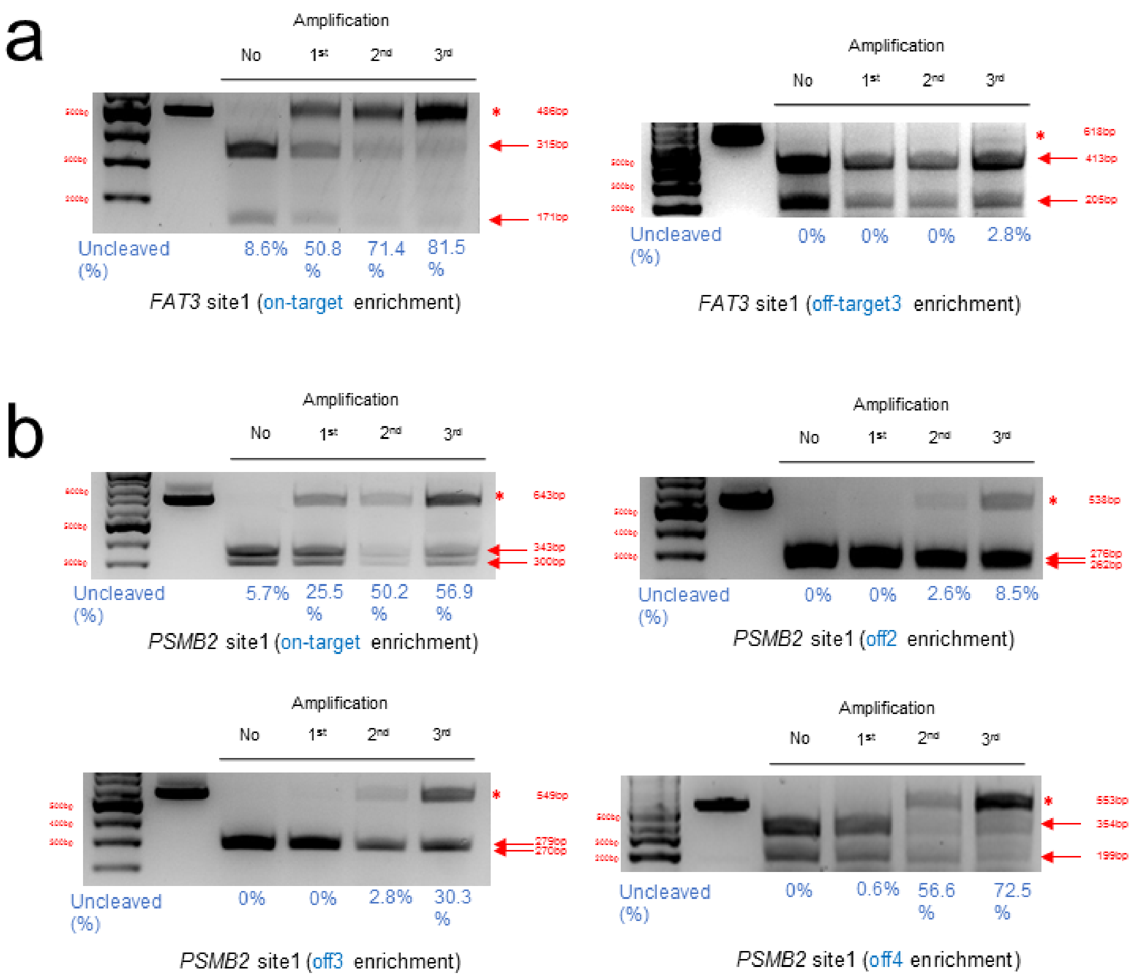


**Supplementary Figure 3. Genotyping of the CRISPR amplification of intracellular off-target mutations induced by CRISPR-Cas9 and adenine base**

**editor(ABE). (a)** The relative mutant DNA frequency(%) for target and predicted Cas9 off-target sites of *FAT3* sequence in genomic DNA from HEK293FT cell is calculated by cleavage and separation of PCR amplicons on 2% agarose gel. **(b)** The relative single-base substituted DNA frequency(%) for target and predicted base editor off-target sites of *PSMB2* sequence in genomic DNA from HEK293FT cell is calculated by cleavage and separation of PCR amplicons on 2% agarose gel. Uncleaved DNA fractions are indicated by asterisk and cleaved DNA fractions are indicated by red arrows.

Supplementary Fig.4

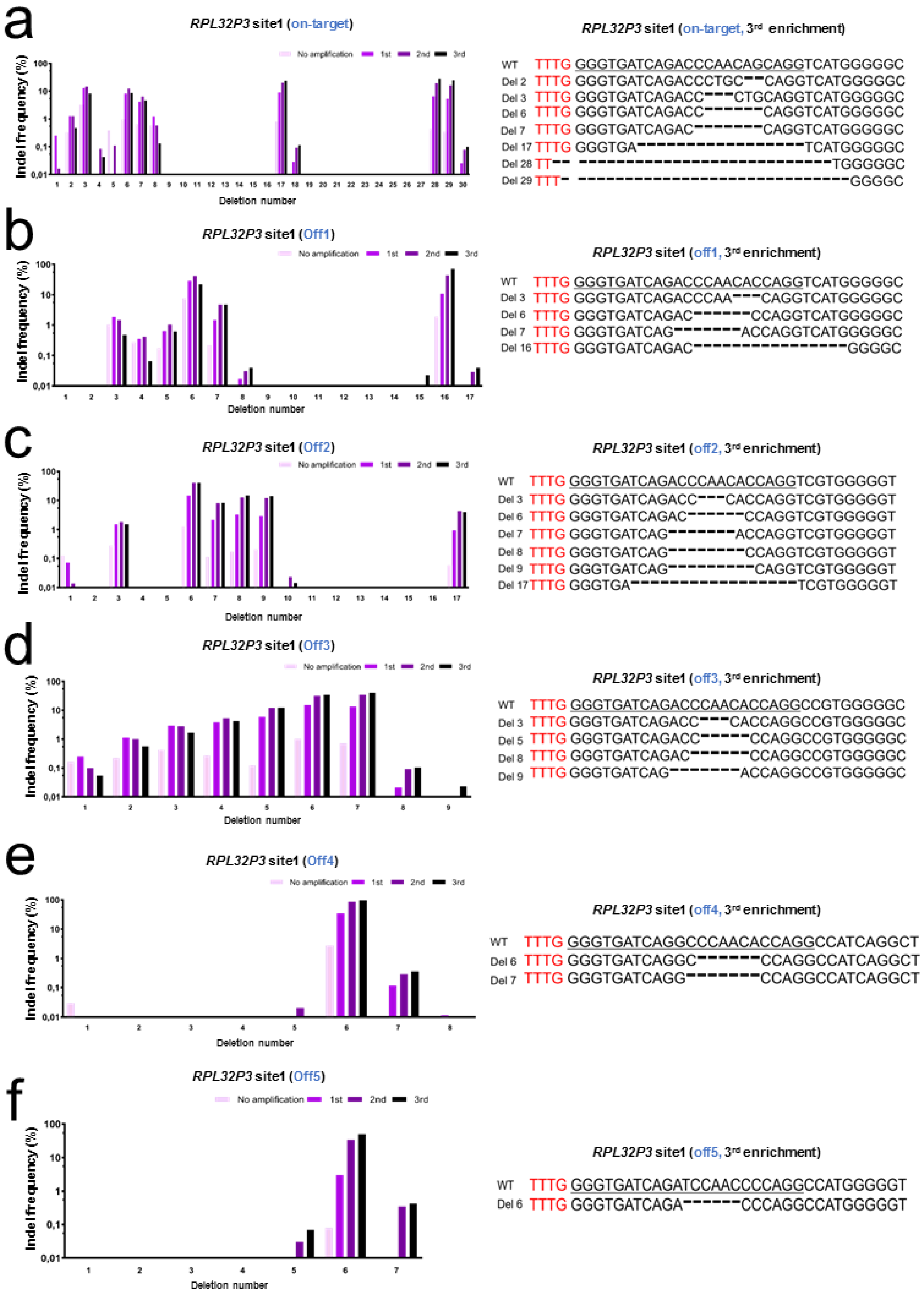


**Supplementary Figure 4. Enriched mutant DNA pattern induced by Cpf1 at predicted off-target sites for RPL32P3 target sequence.** NGS data analysis of

AsCpf1 induced indel patterns enriched by multiple round CRISPR amplification for **(a)** *RPL32P3* on-target site**, (b)** off-target site1, **(c)** off-target site2, **(d)** off-target site3, **(e)** off-target site4, and **(f)** off-target site5, respectively.

Supplementary Fig.5

a

***DNMT1* site3 (On-target)**

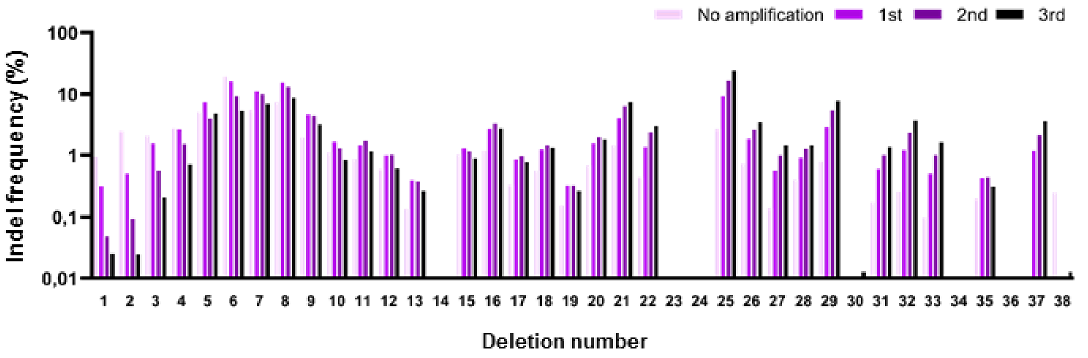


***DNMT1* site3 (On-target, $3^{rd}$ enrichment)**

```
WT ATGTTTC CTGATGGTCCATGTCTGTTACTCGCCTGTCAAGTGGCGT
5  ATGTTTC CTGATGGTCCATG-----TACTCGCCTGTCAAGTGGCGT
6  ATGTTTC CTGATGGTCCATG------ACTCGCCTGTCAAGTGGCGT
7  ATGTTTC CTGATGGTCCA-------TACTCGCCTGTCAAGTGGCGT
8  ATGTTTC CTGATGGTCCAT--------CTCGCCTGTCAAGTGGCGT
9  ATGTTTC CTGATGGTCCATG---------CGCCTGTCAAGTGGCGT
16 ATGTTTC CTGATGGT----------------CCTGTCAAGTGGCGT
21 ATGTTTC CTGATG---------------------GTCAAGTGGCGT
22 ATGTTTC CTGA----------------------TGTCAAGTGGCGT
25 ATGTTT- ------------------------CCTGTCAAGTGGCGT
26 ATGTT-- ------------------------CCTGTCAAGTGGCGT
27 ATGTTTC CTGAT---------------------------GTGGCGT
28 ATGTTTC CTGATGGT----------------------------CGT
29 ATGTTTC CTGA-----------------------------TGGCGT
31 ATGTTTC CTGATG-------------------------------GT
32 A------ --------------------------TGTCAAGTGGCGT
33 ATGT--- ------------------------------AAGTGGCGT
37 ATGTTTC C-------------------------------------T
```

Supplementary Fig.5

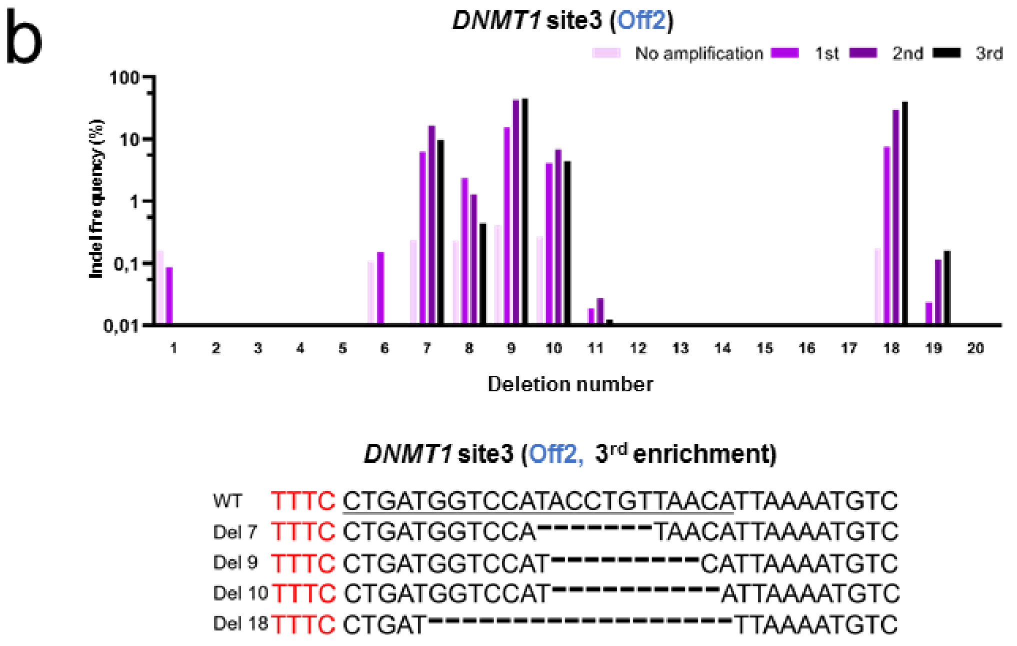


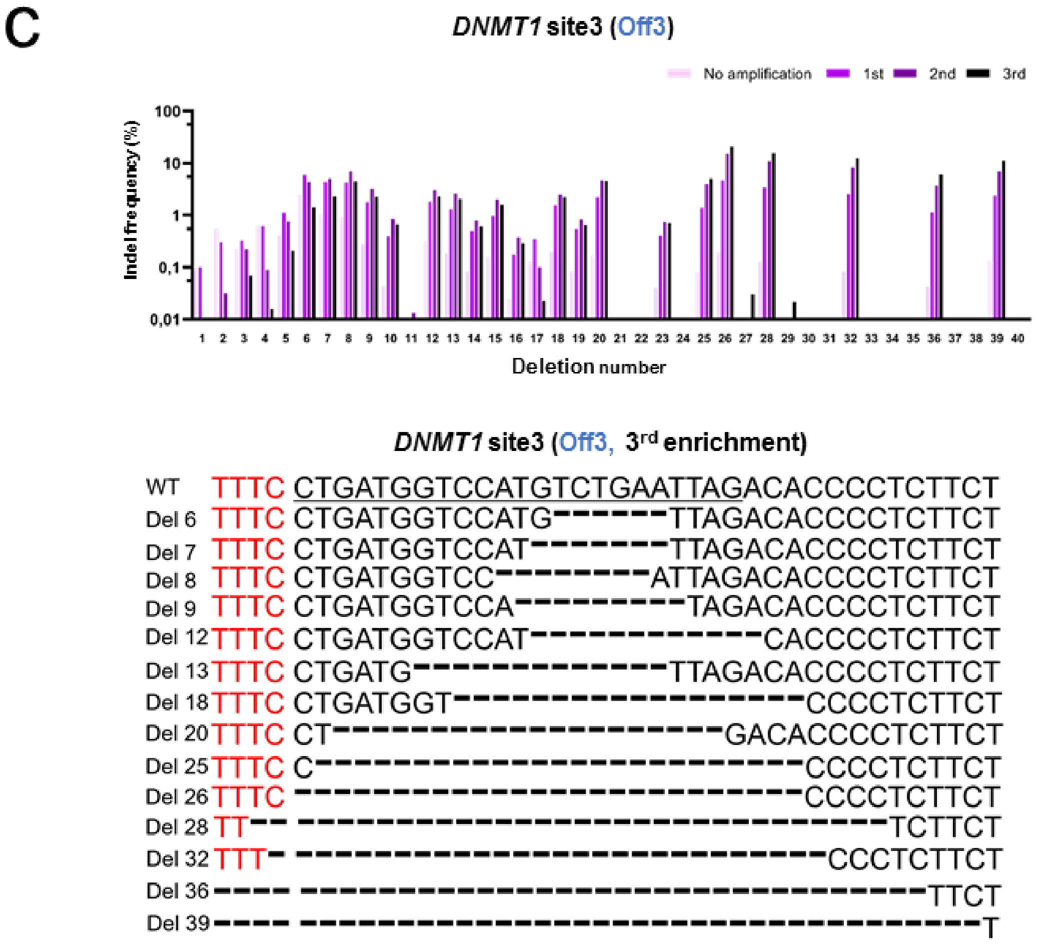


**Supplementary Figure 5. Enriched mutant DNA pattern induced by Cpf1 at predicted off-target sites for *DNMT1* target sequence.** NGS analysis of AsCpf1

induced indel patterns on *DNMT1*-site3 locus enriched by third round CRISPR amplification for **(a)** on-target**, (b)** off-target site2**, (c)** off-target site3, respectively.

Supplementary Fig.6

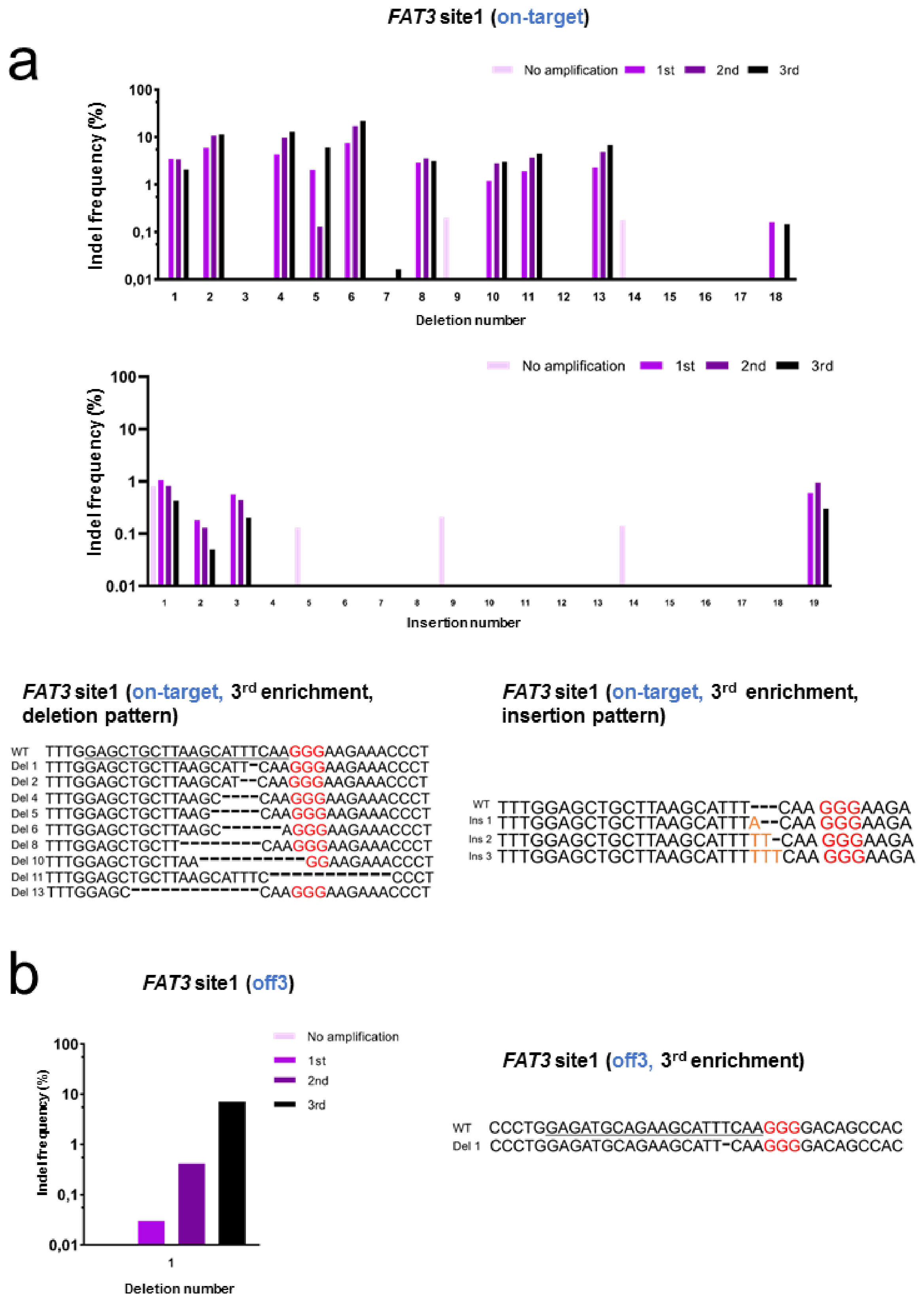


**Supplementary Figure 6. Enriched mutant DNA pattern induced by Cas9 effector at predicted off-target sites for *FAT3* target sequence.** NGS analysis of

SpCas9 induced indel patterns on *FAT3* locus enriched by third round CRISPR amplification for **(a)** on-target and **(b)** off-target site3. PAM sequence(NGG) for SpCas9 is shown in red color and inserted DNA bases are shown in orange color respectively.

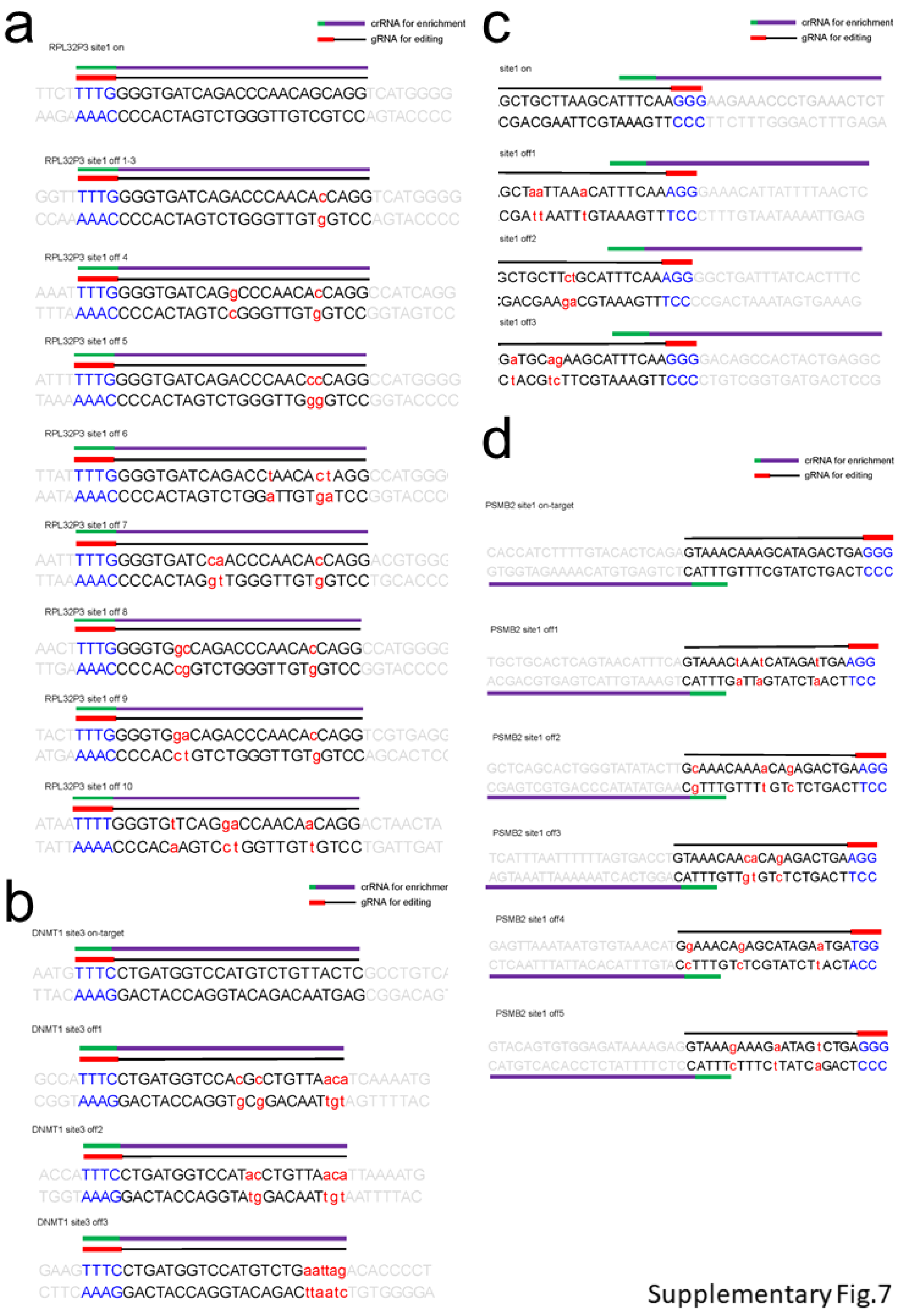


**Supplementary Figure 7. Design of the guide RNA for target specific cleavage and mutant DNA amplification. Guide RNA was used to induce target genomic**

**locus mutation by various effectors and crRNA was used for target DNA enrichment by CRISPR amplification. Each guide and crRNA was designed for (a)** *RPL32P3* on/off site mutation by Cpf1, **(b)** *DNMT1*-site3 on/off site mutation by Cpf1, **(c)** *FAT3* on/off site mutation by Cas9, **(d)** *PSMB2* on/off site mutation by adenine base editor. PAM sequence and mismatched sequences within protospacer region is shown in blue and red color, respectively.
